# Spontaneous fluctuations in global connectivity reflect transitions between states of high and low prediction error

**DOI:** 10.1101/2025.03.18.643969

**Authors:** Paul C. Bogdan, Shenyang Huang, Lifu Deng, Simon W. Davis, Roberto Cabeza

## Abstract

While numerous researchers claim that the minimization of prediction error (PE) is a general force underlying most brain functions, others argue instead that PE minimization drives low-level, sensory-related neuronal computations but not high-order, abstract cognitive operations. We investigated this issue using behavioral, fMRI, and EEG data. Studies 1A/1B examined semantic- and reward-processing PE using task-fMRI, yielding converging evidence of PE’s global effects on large-scale connectivity: high-PE states broadly upregulated ventral-dorsal connectivity, and low-PE states upregulated posterior-anterior connectivity. Investigating whether these global patterns characterize cognition generally, Studies 2A/2B used resting-state fMRI and showed that individuals continuously fluctuate between ventral-dorsal (high-PE) and posterior-anterior (low-PE) dynamic connectivity states. Additionally, individual differences in PE task responses track differences in resting-state fluctuations, further endorsing that these fluctuations represent PE minimization at rest. Finally, Study 3 combined fMRI and EEG data, and the study found that the fMRI fluctuation amplitude correlates most strongly with EEG power at 3–6 Hz, consistent with the PE network fluctuations occurring at Delta/Theta oscillation speeds. This whole-brain layout and timeline together are consistent with high/low-PE fluctuations playing a role in integrative and general sub-second cognitive operations.

## 1. Introduction

The neuroscientific literature is replete with claims that one of the brain’s primary functions is to model its environment, generate predictions, and refine its activity to minimize prediction error (PE). It is generally agreed that the brain uses PE mechanisms at neuronal or regional levels,^1,2^ and this idea has been useful in various low-level functional domains, including early vision^1^ and dopaminergic reward processing.^3^ Some theorists have further argued that PE propagates through perceptual pathways and can elicit downstream cognitive processes to minimize PE.^4–7^ For example, PE might be minimized via attentional re-orienting in parietal regions,^8,9^ memory encoding in the medial temporal lobes,^10,11^ and language learning or category-updating in temporal areas.^12–14^ More controversially, it has also been argued that abstract functions like cognitive processing of emotion,^4^ Theory of Mind,^5^ and others^7,15^ have likewise been described as PE minimization. Following this line of reasoning, even cognition without exogenous stimuli may reflect the minimization of PE. For instance, daydreaming about what food you will eat later can be seen as an effort to avoid future sub-optimal food choices. Coordinating cortical mechanisms for sensation, decision-making, and autobiographical memory, one can describe cognition as a series of mental operations seeking to minimize PE.

PE minimization may broadly coordinate brain functions of all sorts, including abstract cognitive functions. This includes the types of cognitive processes at play even in the absence of stimuli (e.g., while daydreaming). While it may seem counterintuitive to associate this type of cognition with PE – a concept often tied to external surprises – it has been proposed that the brain’s internal generative model is continuously active.^16–18^ Spontaneous thought, such as planning a future event or replaying a memory, is not a passive, low-PE process. Rather, it can be seen as a dynamic cycle of generating and resolving internal uncertainty. While daydreaming, you may be reminded of a past conversation, where you wish you had said something different. This situation contains uncertainty about what would have been the best thing to say. Wondering about what you wish you said can be viewed as resolving this uncertainty, in principle, forming a plan if the same situation ever arises again in the future. Each iteration of the simulated conversation repeatedly sparks and then resolves this type of uncertainty.

Ideas about PE operating at a global level and underlying abstract cognitive functions are difficult to evaluate experimentally, which is the root of the controversy about this view.^19–21^ In particular, it is challenging to isolate high-level and multifaceted cognition processes in a way that deters alternative explanations linked to other neural mechanisms or stimulus-specific functions. Yet, doing so would yield key evidence and provide direction for the variety of neurocognitive theories attempting to unify all levels of brain function under a PE lens.^4–7,15^ We pursued this goal using three studies (**Figure 1**) that collectively targeted a specific question: Do the task-defined connectivity signatures of high vs. low PE also recur during rest, and if so, how does the brain transition between exhibiting high/low signatures?

**Figure 1.**
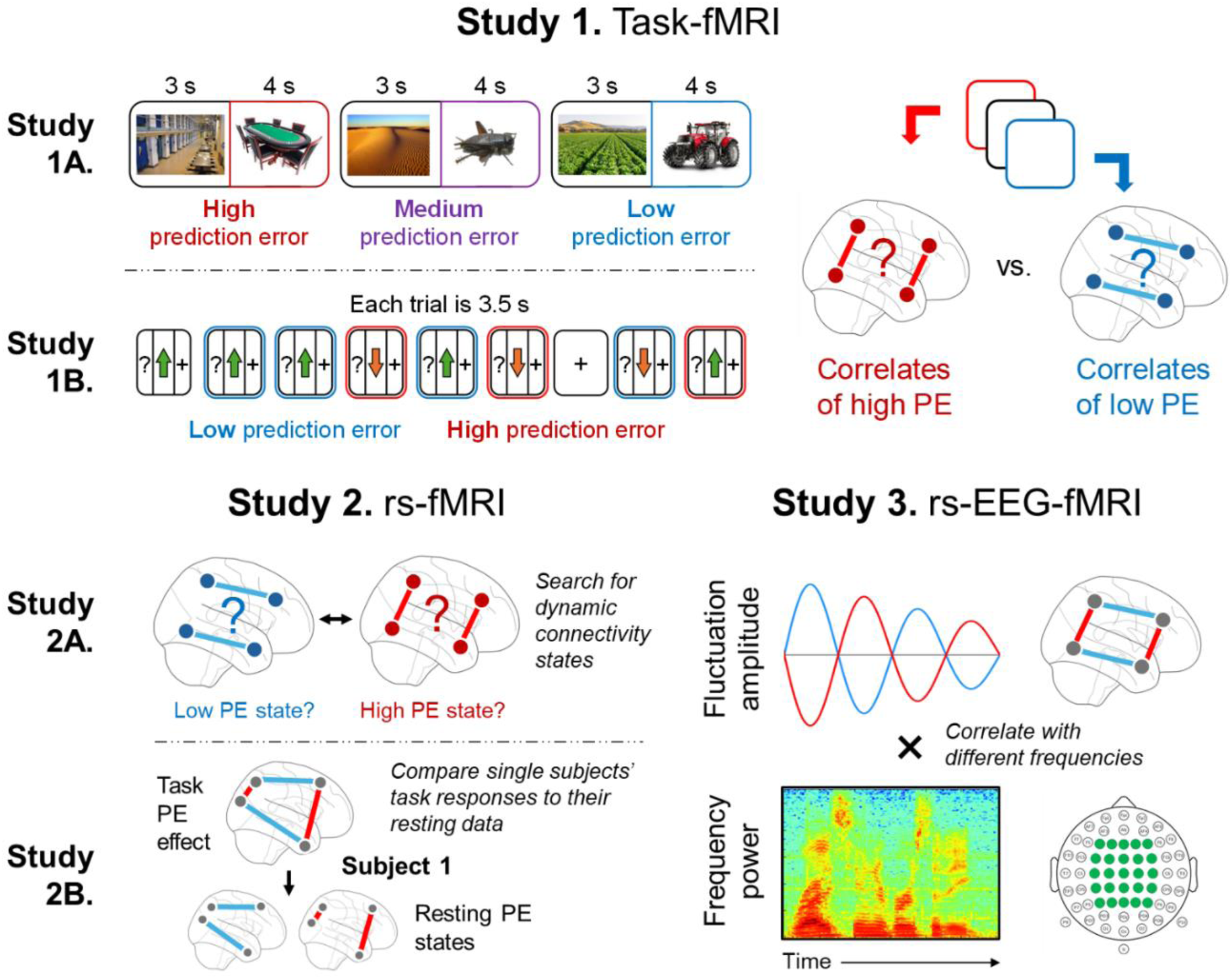
Task design and graphical summaries of analytic strategies. (Study 1) Two task-fMRI studies were conducted to identify the connectivity patterns upregulated by high or low PE. **(A)** In each trial, participants saw a scene-object pair and were asked to report how likely it would be to find the object in the scene using a 4-point scale that was active while the object was on screen. **(B)** Separate participants completed a gambling task wherein each trial they made a binary choice (“?” screen) and were then told whether the choice caused them to earn money (upward arrow) or lose money (downward arrow). The “+” panel represents a 15 s fixation period between blocks. Both earning or losing money could elicit high or low PE, depending on whether it deviates from prior trials’ outcomes. **(Study 2) (A)** Using rs-fMRI data collected from Study 1A participants, analyses examined changes in connectivity over time to assess whether participants spontaneously fluctuate between the high and low PE signatures found in Study 1. **(B)** Further analyses examined whether slight differences in Study 1B participants’ task-fMRI responses to PE mirrored differences in their rs-fMRI connectivity states. **(Study 3)** With rs-EEG-fMRI data from another pool of participants, analyses modeled the amplitudes of fMRI connectivity fluctuations over time and correlated this with EEG oscillatory power at different frequencies to assess which frequency best tracked the fMRI patterns.

First, we used task-fMRI to identify the functional connectivity signatures of high-PE and low-PE states. Few prior PE studies have targeted connectivity, contrasting the hundreds that have targeted BOLD.^22,23^ Yet, compared to measuring activity, tracking connectivity better taps into the core concepts of the integrative PE view that theorizes how signals communicate through the brain to propagate and/or minimize PE. For Study 1A, we conducted a task-fMRI study (N = 66) that elicited PE relative to pre-existing schemas and aimed to identify the connectivity patterns of low and high PE. For Study 1B, we analyzed a public task-fMRI study (N = 1000) that elicited reward PE using a gambling task. We expected to replicate the connectivity signatures of high and low PE seen in the schema task, showing that these networks emerge across starkly different PE tasks.

Second, we reasoned that if PE processing characterizes cognition more generally, then it should also occur in the absence of sensory stimuli. Studies 2A and 2B examined PE during resting-state fMRI (rs-fMRI) scans. Detecting PE processing during a resting-state fMRI scan required a template of PE processing.^24^ Thus, we used the connectivity patterns of high and low PE identified in Studies 1A/1B, and searched the rs-fMRI data for these connectivity patterns. This strategy can link the Study 1 task manipulations and resting-state phenomena without *a priori* assumptions about individual brain regions’ exact roles.^25^ One potential problem of this approach is known as “reverse inference”: Finding the same networks during a PE task and during rest does not mean that the networks’ engagement during rest necessarily reflects PE processing. To engage with this issue, we examined if the engagement of putative PE networks during rest and during a PE task was significantly correlated across participants. Although this does not entirely address the reverse inference dilemma and can only produce correlative evidence, the present research nonetheless investigates these widely speculated upon PE ideas more directly than any prior work.

Finally, in Study 3, we used rs-fMRI-EEG simultaneous recordings (N = 21) to investigate whether participants fluctuate between high/low PE states at speeds consistent with whole-brain integrative cognitive operations. Although no prior fMRI study has focused on PE under this lens, this perspective is motivated by EEG research. Specifically, in EEG studies on executive function, PE detection elicits a frontal negativity around 100-400 ms^26,27^ that is followed by a centroparietal positivity around 300-800, indexing PE minimization processes such as redirecting attention, changing behavior, and updating working memory.^26,28,29^ These results indicate that shifts from high-to low-PE states can occur within a few hundred milliseconds. This idea is consistent with evidence that PE modulates Delta (1-4 Hz) and Theta (4–8 Hz) power, which aligns with the timing of error-related negativity/positivity.^30^ Hence, in Study 3, we expected to find that participants’ high- and low-PE networks would fluctuate multiple times per second. Although such speed outpaces the temporal resolution of fMRI, correlating fluctuations in dynamic connectivity measured from fMRI data with EEG oscillations can provide an estimate of the fluctuations’ speed. This interpretation of a correlation again runs up against issues related to reverse inference but would nonetheless serve as initial suggestive evidence that spontaneous transitions between network states occur rapidly.

Instead, sub-second fluctuations could reflect individual cognitive operations, such as working memory updating, attentional shifting, or other processes attributed to PE minimization. Moreover, we specifically expected that the fluctuations occur at Delta/Theta frequencies, which would also track the PE literature on oscillations and further validate that the resting-state patterns reflect PE minimization^30^ – that is, despite concerns about reverse inference and interpreting rs-fMRI data, by demonstrating an additional link to Delta/Theta, this would constitute suggestive converging evidence for the PE interpretation

In sum, we investigated the global and task-general/spontaneous role of PE in three steps. First, we used task-fMRI to identify large-scale brain networks associated with high and low PE. Second, we examined the recruitment of these networks during rs-fMRI, and although the problems related to reverse inference are impossible to overcome fully, we engage with this issue by linking rs-fMRI data directly to task-fMRI data of the same participants, which can provide suggestive evidence that the same neural mechanisms are at play in both. Finally, we examined rs-fMRI-EEG data to assess whether we find parallels consistent with the high/low-PE network fluctuations occurring at fast timescales suitable for the type of cognitive operations typically targeted by PE theories.

## 2. Results

### 2.1. Study 1 – Task-fMRI: Mapping the effects of prediction error on connectivity

#### 2.1.1. Study 1A – Manipulating semantic PE

Participants completed a task wherein each trial they viewed a well-known scene, such as a farm. The scene was followed by either an object that was highly predictable given the scene (low PE; e.g. a tractor in a farm), an object that was less predictable but still possible (medium PE; e.g., an insect in a desert), or an object that was highly unpredictable (high PE; e.g. a poker table in a prison). Participants rated how likely it would be for each object to appear in its associated scene (1 = “*Very unlikely*”, 4 = “*Very likely*”), which confirmed that participants processed the stimuli as intended: Low-PE objects were rated as likely to appear in the preceding scene (M = 3.67, SD = 0.20), medium-PE objects as moderately likely (M = 2.11, SD = 0.30), and high-PE objects as unlikely (M = 1.28, SD = 0.24). Average response times for trials where participants gave the expected response were slightly slower for the high-PE condition (M = 1.87 s, SD = 0.40 s) compared to the low-PE condition (M = 1.79 s, SD = 0.41 s; *t*[65] = 2.54, *p* = .014, *d =* 0.27). This response time gap is consistent with error-related slowing, although it is nominally small.^31^ Overall, these behavioral trends suggest the suitability of the data for understanding the neural correlates of PE.

In the fMRI data, we compared the three PE conditions’ beta-series functional connectivity, aiming to identify network-level signatures of PE processing, from low to high. Network-level signatures could later be leveraged for detecting global trends in rs-fMRI data. After demonstrating that PE significantly influences brain-wide connectivity using Network-Based Statistic analysis (Supplemental Materials 2.1)^32^, we conducted a modularity analysis to study how specific groups of edges are all sensitive to high/low-PE information. For the modularity analysis, we first defined a connectome matrix of beta values, wherein each edge’s value was the slope of a regression predicting that edge’s strength from PE (coded as Low = -1, Medium = 0, High = +1; **Figure 2A**). Modules representing positive and negative effects were then identified by applying a variant of Louvain’s algorithm twice: once for a binary matrix of the 5% of edges showing the strongest Low > High effects and once for the analogous High > Low matrix (**Figure 2B**).

**Figure 2.**
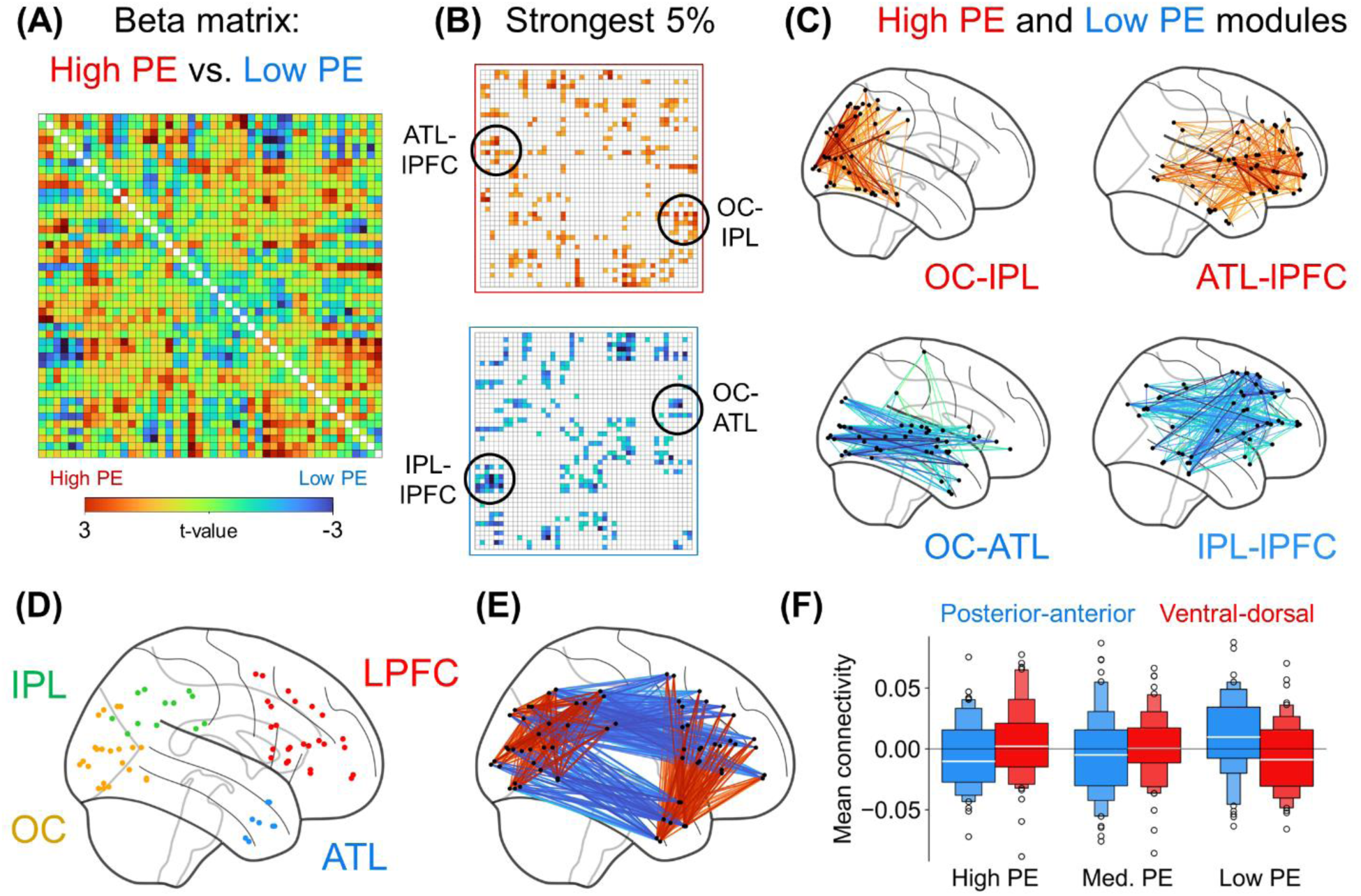
Modularity behind the enhancing and weakening effects of prediction error. **(A)** Each neocortical connection was submitted to a regression predicting connectivity strength based on PE. For clearer visualization, the shown 46 x 46 matrix simplifies the original 210 x 210 matrix by averaging ROIs per anatomical labels. Supplemental Figure S1 shows the exact ROIs used on a glass brain, illustrating their precise anatomy. **(B)** Modularity was assessed in terms of the 5% of edges showing the most positive coefficients and 5% showing the most negative coefficients. **(C)** Two high-PE modules and two low-PE modules are shown. **(D & E)** The four modules can be described anatomically as connections among four anatomical quadrants. **(F)** Analyzing these anatomically defined connections shows a significant Connectivity-Direction × PE interaction (β [standardized] = .20, p < .001). Boxes indicate quantiles.

Roughly, the high/low PE modules appear as connections among quadrants, consisting of the occipital cortex (OC), inferior parietal lobule (IPL), anterior temporal lobe (ATL), and dorsolateral prefrontal cortex (LPFC, i.e., the inferior frontal gyrus [IFG] & middle frontal gyrus [MFG]) (**Figure 2C-2E**). That is, high PE enhanced ventral-dorsal connectivity (OC-IPL & ATL-LPFC) whereas low PE enhanced posterior-anterior connectivity (OC-ATL & IPL-LPFC) (**Figure 2F**). This four-quadrant description omits some ROIs included in the modules (e.g., the lateral temporal component of the IPL-lPFC module; **Figure 2C**).

Some ROIs appear in **Figure 2C** but are excluded from the four targeted quadrants (**Figures 2C & 2D)** – e.g., posterior inferior temporal lobe and fusiform ROIs are excluded from the OC-IPL module, and middle temporal gyrus ROIs are excluded from the IPL-LPFC modules. These exclusions, in favor of a four-quadrant interpretation, are motivated by existing knowledge of prominent structural pathways among these quadrants. This interpretation is also supported by classifier-based analyses showing connectivity within each quadrant is significantly influenced by PE (Supplemental Materials 2.2), along with analyses of single-region activity showing that these areas also respond to PE independently (Supplemental Materials 3). Hence, we proceeded with further analyses of these quadrants’ connections, which summarize PE’s global brain effects.

This four-quadrant setup also imparts analytical benefits. First, this simplified structure may better generalize across PE tasks, and Study 1B would aim to replicate these results with a different design. Second, the four quadrants mean that each ROI contributes to both the posterior-anterior and ventral-dorsal modules, which would benefit later analyses and rules out confounds such as PE eliciting increased/decreased connectivity between an ROI and the rest of the brain. An additional, less key benefit is that this setup allows more easily evaluating whether the same phenomena arise using a different atlas (Supplemental Materials 7).

#### 2.1.2. Study 1B – Manipulating reward PE

To confirm that the ventral-dorsal and posterior-anterior networks are indeed general signatures of high and low PE, respectively, we investigated their occurrence during a very different PE task: the gambling task from the Human Connectome Project (N = 1000).^33,34^ In each trial of this task, participants made a binary choice, then won or lost money as pre-determined. Our analyses modeled PE as the absolute difference between a trial’s monetary outcome and participants’ expectations given recent prior trials (i.e., relative to an exponential moving average). This strategy assigns a continuous PE measure to each trial, which was then split at the median to define high/low PE conditions. As a validation check, we confirmed that participants’ response times were slower after loss trials with high-PE feedback (M = 422 ms) compared to after loss trials with low-PE feedback (M = 418 ms) (*t*[999] = 7.46, p < .0001). This is consistent with expected post-error/surprise slowing,^31^ although given how slight the response-time difference was (4 ms), it was not expected to reflect any meaningful confound.

Analyses evaluated the consistency between the connectivity effects of PE in the present gambling task and those seen earlier in the semantic PE task. Here, functional connectivity was computed for high/low PE conditions. Then, a paired t-test between the two conditions was performed for each edge, which yielded a connectome-wide high/low contrast. This contrast matrix paralleled that of Study 1A, as evidenced by element-wise correlations of the matrices’ bottom triangles (**Figures 2A & 3A**, *r* = .27, *p* < .0001). Critically, the present data also reproduced Study 1A’s Connectivity Direction × PE interaction (*F*[1, 999] = 68.41, *p* < .0001; **Figure 3B**). This prominent correlation and reproduction point to strong task-general qualities of the identified global PE phenomena.

**Figure 3.**
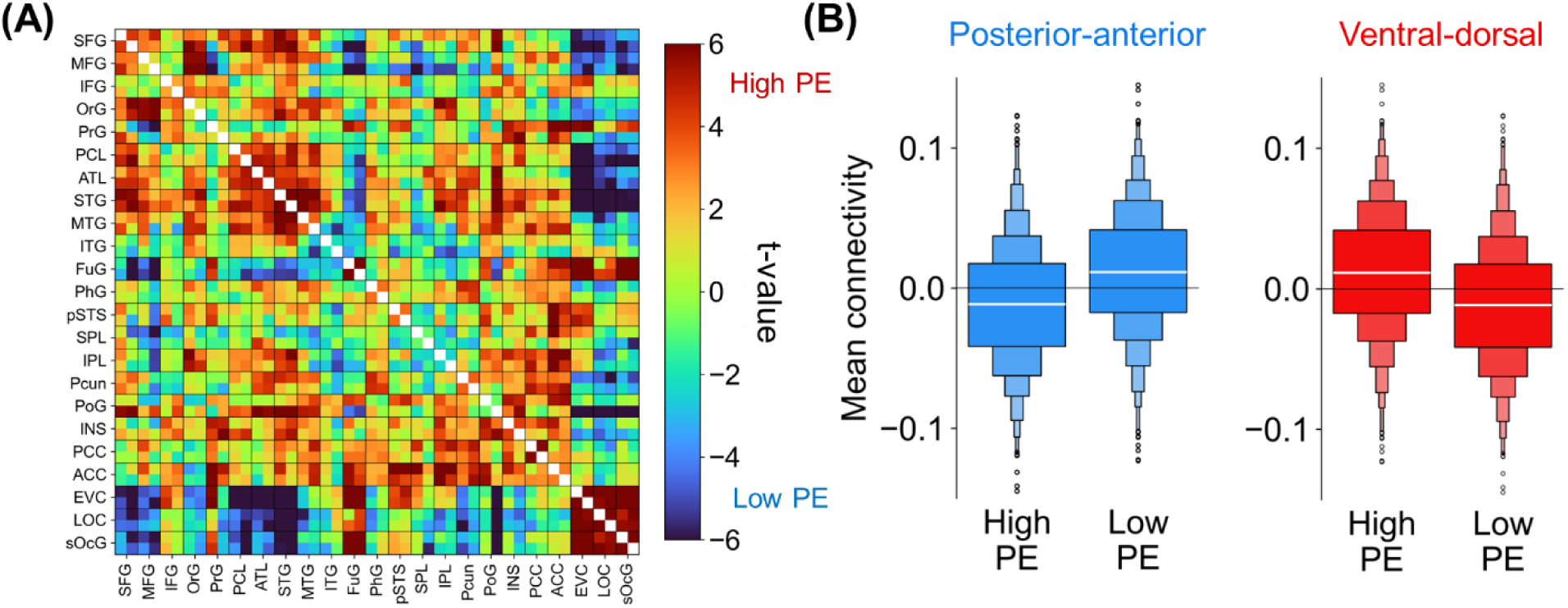
Reproducing the posterior-anterior/ventral-dorsal PE effects using a gambling task. **(A)** This matrix represents high/low PE paired t-tests for each connection. Like Figure 2, a 46 x 46 regional matrix is shown rather than a 210 x 210 ROI matrix for clarity. **(B)** Low PE enhances posterior-anterior connectivity while high PE enhances ventral-dorsal connectivity but note the High > Low PE ventral-dorsal effect is driven by ATL-LPFC connectivity, and the OC-IPL effect is minuscule. For the bars, the connectivity data was normalized by subtracting means across conditions. Boxes indicate quantiles.

In sum, Study 1A identified distinct global connectivity patterns linked to PE: High PE upregulates a ventral-dorsal network whereas low PE upregulates a posterior-anterior network. Study 1B reproduced these signatures using a very different PE task. This regularity points to PE processing as prompting general connectome-wide mechanisms. Yet, it remains to be tested whether PE processing applies to high-level cognition more broadly, including in the absence of stimuli specifically designed to elicit PE. Thus, Study 2 investigated whether the ventral-dorsal and posterior-anterior networks are engaged when stimuli are absent, as would be expected if PE minimization is a fundamental function of the human brain.

### 2.2. Study 2 – Rs-fMRI: Identifying connectivity fluctuations tied to prediction error

#### 2.2.1. Study 2A – Dynamic connectivity and global states

We assessed whether the above correlates of high/low PE emerge as dynamic connectivity states in rs-fMRI data from Study 1A participants. In other words, we essentially asked whether resting-state participants are sometimes in low PE states and sometimes in high PE states, which would be consistent with spontaneous PE processing in the absence of stimuli. To formally test this, we first measured *time-varying connectivity* on a TR-by-TR basis.^35^ For example, OC-IPL connectivity strength in the first TR is the product of OC and IPL standardized activations in said TR (OC_BOLD_ × IPL_BOLD_). Time-varying connectivity was computed for the edges among adjacent portions of the earlier four quadrants (**Figure 4A**). In turn, edge-edge correlations^35^ were calculated to form the **Figure 4B** matrix; this matrix shows, for example, that strong L OC-IPL connectivity coincides with weak L OC-ATL connectivity. Submitting this matrix to Louvain’s algorithm, in turn, yielded the two modules in **Figure 4C**: Here, module 1 indicates that, when IPL-LPFC connectivity is strong, then OC-ATL tends to also be strong, and these two sets of edges make up a global state of strong posterior-anterior connectivity. Likewise, module 2 shows that OC-IPL and ATL-LPFC connectivity make up a ventral-dorsal state. These emerging states overlap strikingly with the previous task effects of PE, suggesting that rs-fMRI scans exhibit fluctuations that resemble the signatures of low- and high-PE states.

**Figure 4.**
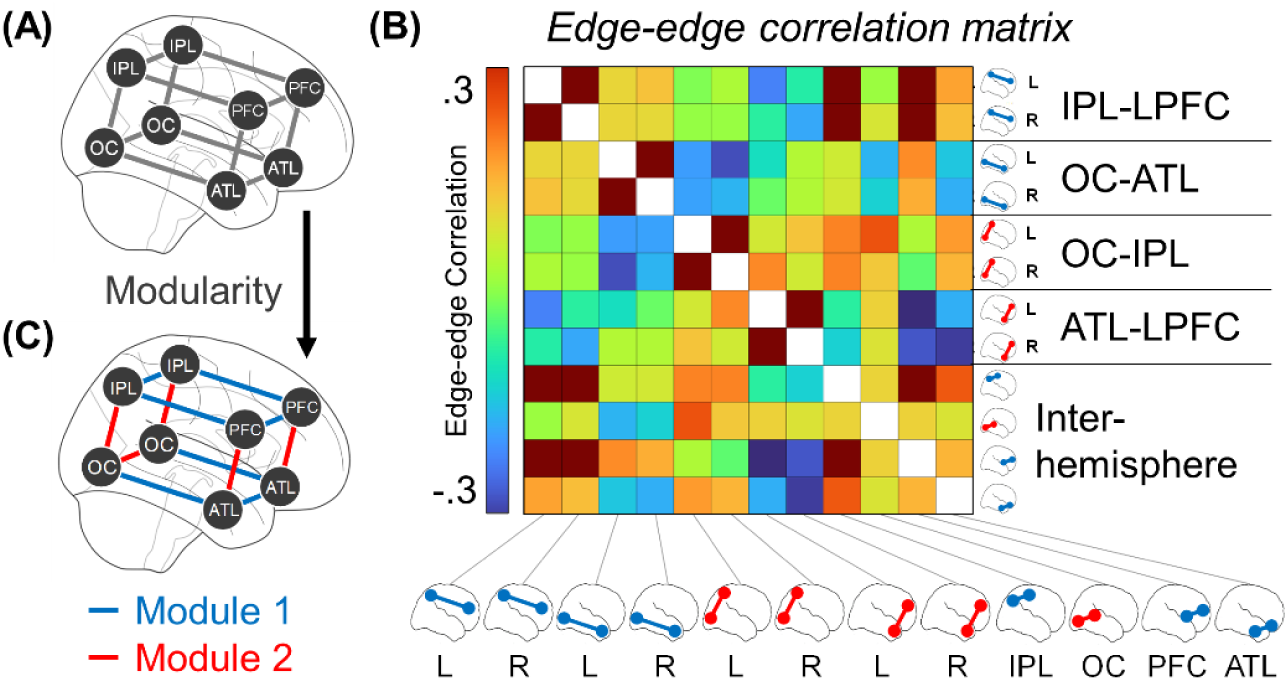
Dynamic connectivity modules. **(A)** Time-varying connectivity was computed between adjacent portions of the four bilateral quadrants from earlier. Given that each quadrant is composed of many ROIs, this involved computing time-varying connectivity for individual ROI-ROI pairs and averaging them to produce overall estimates. **(B)** Based on the time-varying connectivity of each edge, an edge-edge correlation matrix was produced. **(C)** Computing modularity of the edge-edge correlation matrix yielded the two illustrated modules.

#### 2.2.2. Study 2B – Linking task and resting-state PE

Study 2B had two goals. The first goal was to replicate the results of Study 2A in a different group of rs-fMRI participants (N = 1000). Fulfilling this goal, Study 2B yielded the same posterior-anterior and ventral-dorsal connectivity patterns during rs-fMRI (**Figure 5A**).

**Figure 5.**
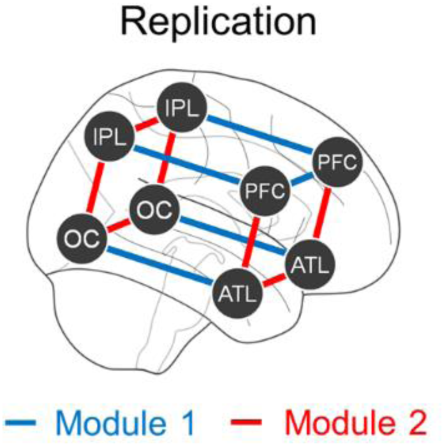
Replication of the connectivity modules and subject-specific analyses. Performing the Study 2A modularity analysis using the Study 1B participants yields a similar pair of modules as in Figure 4 but with slight changes to the interhemispheric connections.

The second goal of Study 2B was to engage with the aforementioned reverse inference issue: Finding the same networks during a PE task and during rest does not imply that the networks’ engagement during rest necessarily reflects PE processing. By establishing these networks using two very different PE tasks (Studies 1A/1B), this deters non-PE interpretations of the networks specific to just one task. To further address this issue, we considered the evidence that the anatomical organization of functional networks varies between individuals.^36^ Hence, if the recruitment of ventral-dorsal/posterior-anterior networks during rest is indeed related to PE, then a participant’s unique connectivity response to PE in the reward-processing task should match their specific patterns of resting-state fluctuation. Our focus is not on the overall strength of the task contrast and overall fluctuation magnitude at rest; the theoretical justification for these being linked seems questionable, as PE minimization is presumed to characterize every cognitive process of every single person’s brain. Yet, there should still be slight anatomical differences in where exactly these networks are positioned, and these differences should be consistent across the task-fMRI and rs-fMRI data. We tested whether this was indeed the case.

To evaluate anatomical correspondence across task-fMRI and rs-fMRI, we analyzed these effects for subsets of the quadrants. From each quadrant, we randomly selected a pair of bilateral ROIs (one L ROI and its symmetric R ROI). Then, using just these eight ROIs, we calculated the strength of each participant’s task-fMRI effect (Connectivity Direction × PE interaction) along with the magnitude of their rs-fMRI posterior-anterior/ventral-dorsal fluctuation (mean absolute difference of time-varying connectivity, E[|PA – VD|]; see Study 3 and Methods). These two measures were computed for all 3,432 possible sets of eight ROIs. For each participant, the within-subject correlation was computed with respect to the 3,432 ROI sets with the expectation that these would be positive on average. A positive correlation would indicate that ROIs associated with stronger PE effects in the task are linked to stronger fluctuation at rest. This was indeed found and demonstrated to be significantly above zero by submitting the 1000 participants’ correlations to a one-sample t-test (mean *r* = .021, *t*[999] = 3.31, *p* < .001). Note that this nominally low correlation likely stems from the short task-fMRI scan length (only six minutes), which in turn causes the split-half reliability of a participant’s task contrast to be just .08 (the reliability of the rs-fMRI measure was much higher, see Supplemental Materials 4). Hence, a nominally low correlation is to be expected given that reliability sets an upper limit on the possible correlation. This rs-fMRI effect is specific to the PE contrast in the task-fMRI data; if the same analysis is performed with respect to a win/loss task-fMRI contrast, this produces an insignificant task-fMRI × rs-fMRI association (mean *r* = .006, *t*[999] = 1.18, *p* = .24). Hence, a participant’s unique pattern of connectivity elicited by PE matches their fluctuations in resting-state scans.

To be clear, this does not entirely dissuade concerns about reverse inference, which would require a type of causal manipulation that is difficult (if not impossible) to perform in a resting state scan. Nonetheless, these results provide further evidence consistent with our interpretation that the resting brain spontaneously fluctuates between high/low PE network states.

Studies 2A and 2B show that, even in the absence of stimuli, participants fluctuate between global connectivity states corresponding to high and low PE. Overall, these results endorse claims that abstract cognition is rooted in PE minimization. However, claims of PE unifying psychological concepts generally refer to mental operations that can occur fairly quickly – e.g., emotional responses, logical inferences, memory retrieval, attentional shifting, or working memory updating. This may seem to create a discrepancy with the results thus far, which are all based on fMRI with its slow temporal resolution. Yet, slow measurement does not imply that underlying neural processes are slow. Study 3 investigated this further.

### 2.3. Study 3 – Rs-fMRI-EEG: Measuring fluctuation frequency

To investigate the temporal properties of our putative global PE networks shifting between high/low PE states, we conducted analyses using fMRI-EEG simultaneous recordings from participants at rest. Our overarching strategy for detecting the fluctuation rate involved first modeling the amplitude of the fMRI connectivity fluctuations on a TR-by-TR basis. Then, we correlated this amplitude with EEG oscillatory power at different frequencies to assess which best aligned with the fMRI patterns (**Figures 6A & 6B**). We validated this strategy with simulations (Supplemental Materials 5). In brief, we specifically simulated a neural oscillator that varies in amplitude over time (**Figures 6C-E**) and reflects opposing posterior-anterior (PA) and ventral-dorsal (VD) connectivity (**Figure 6F**). fMRI can measure time-varying connectivity but only at a slow scale (i.e., modeled as the average connectivity across a TR; **Figure 6G**). Nonetheless, the frequency of the oscillator producing these fMRI measurements can be discerned by linking the magnitude of fMRI fluctuations (|PA – VD| in **Figure 6H**) to EEG oscillatory power at different frequencies (**Figures 6I-K**). If VD and PA strengths fluctuate at 3 Hz, then |VD – PA| in fMRI will most strongly correlate with 3 Hz power in EEG. Notice how the dots in **Figure 6H** follow the dots in **Figure 6J** (3 Hz) better than the dots in **Figure 6I** (0.5 Hz) or **Figure 6K** (10 Hz); this visual comparison is intended for illustrative purposes only, and quantitative analyses are provided in Supplemental Materials 5. This same logic can be applied to experimental data to determine the frequency at which individuals fluctuate between posterior-anterior and ventral-dorsal dynamic connectivity states.

**Figure 6.**
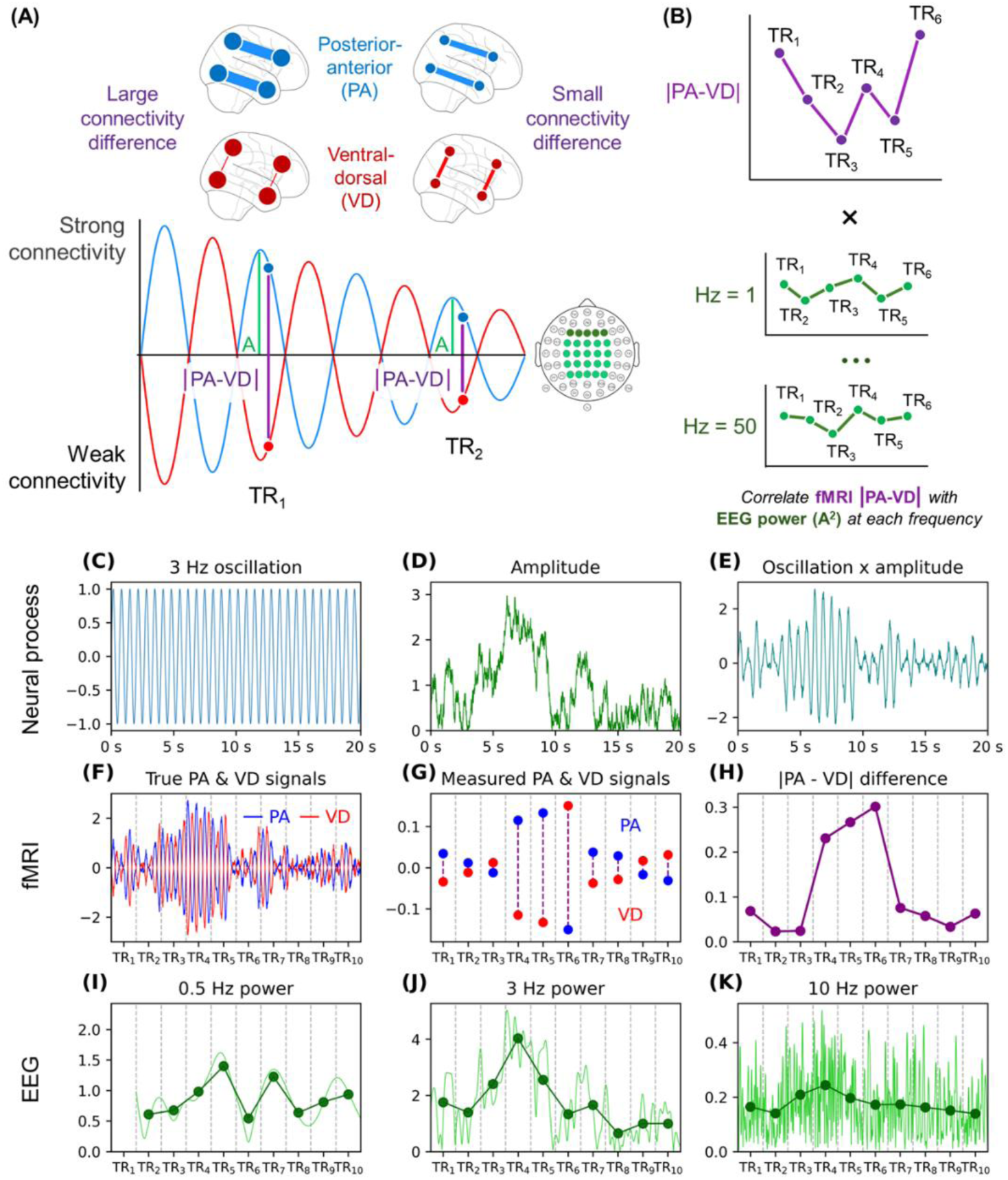
Correlating connectivity fluctuations and oscillatory power. **(A)** If posterior-anterior (PA) and ventral-dorsal (VD) time-varying connectivity measurements at each TR reflect samples from an oscillator, then their absolute difference (|PA – VD|) will correlate with the oscillator’s true amplitude (A) even if the oscillator’s frequency outpaces fMRI’s temporal resolution. **(B)** Correlations between |PA-VD| and amplitude/power at different frequencies allow assessing the frequency of the oscillator generating the PA/VD fluctuations; amplitude (A) and power (A^2^) are equivalent for the analyses, which used Spearman correlations. The fMRI-EEG analytic strategy is motivated by simulations: **(C)** One sine wave is used as a simulation input. **(D)** An autoregressive time series is also used as an input. **(E)** The product of the sine wave and autoregressive time series defined the simulated oscillator, which represents how fluctuations between states may vary in intensity over time. **(F)** Ventral-dorsal (VD) connectivity is defined as the sine-amplitude product, and posterior-anterior (PA) connectivity is defined as the inverse of this product. **(G)** Representing brain data recorded using fMRI, the VD and PA dots here are the average over the TR windows. No hemodynamic response function convolution was applied when making this figure for the sake of clarity, but convolution was performed for the actual simulation results in Supplemental Materials 5. **(H)** Using the PA and VD dots, the |PA – VD| differences were computed. **(I-K)** Green lines represent the time-frequency decomposition of the sine-amplitude product. The dots represent the power averaged for each TR window. The first TR window for 0.5 Hz power was removed as it was susceptible to edge artifacts.

Our analyses used a public dataset of fMRI-EEG simultaneous recordings from participants at rest (N = 21).^37^ We first measured time-varying connectivity for the ventral-dorsal and posterior-anterior edges, as in Study 2B. We calculated the absolute difference between the two at each TR, |PA – VD| to generate connectivity-amplitude time series. Turning to the EEG data, we measured oscillatory power at each frequency from 0.5 to 50 Hz (bin size = 0.5 Hz). To match the power time series with fMRI’s temporal resolution, the power data were averaged by TR (e.g., averaging 0-2099 ms to align with the first 2.1 s TR volume) and convolved with the hemodynamic response function.^38^

Spearman correlations were measured between the connectivity-amplitude time series and each frequency’s power time series. Group statistics and effect sizes were then established by submitting each participant’s correlation to a one-sample t-test relative to zero. The analysis was first done using power averaged across frontal electrodes, as these are the typical focus of PE research on oscillations.^30,39–41^ Correlations revealed that connectivity-fluctuation amplitude was most associated with frontal Delta/Theta and specifically 3-6 Hz power (**Figure 7A**, **Table 1**).

**Figure 7.**
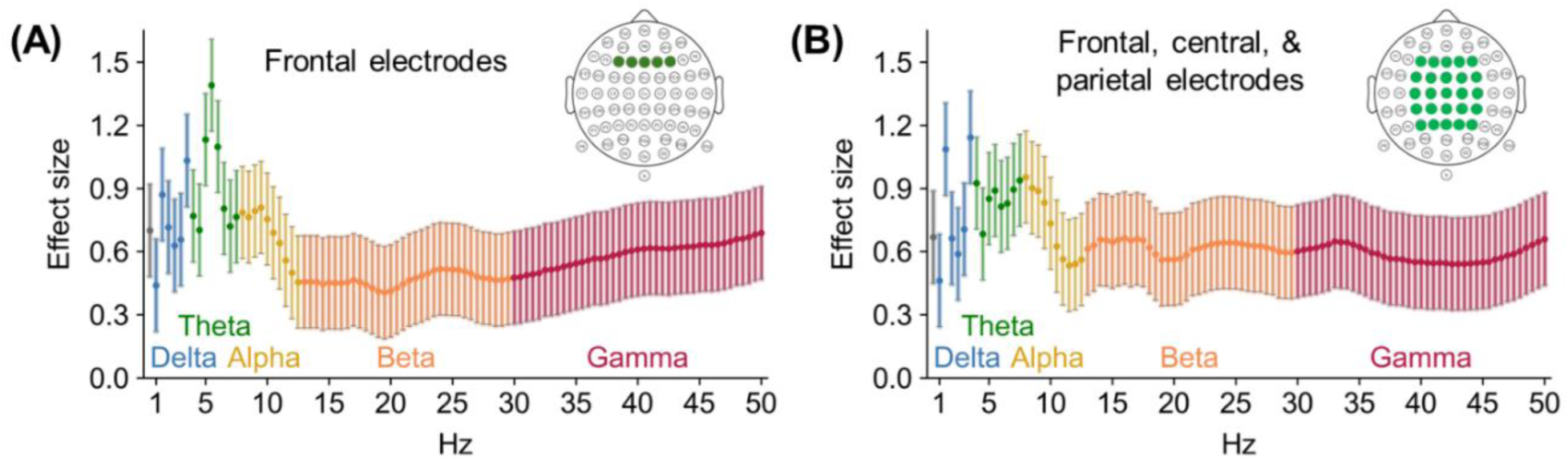
Correlations between fluctuation amplitude and oscillatory power. For all participants, Spearman correlations were measured between connectivity-fluctuation amplitude and power at every 0.5 Hz interval from 0-50 Hz. Visualized here, each dot represents the Cohen’s d effect size from a one-sample t-test assessing whether the mean correlation surpassed zero. This was done using either **(A)** power averaged across the shown frontal electrodes or **(B)** power averaged across the shown frontal/central/ parietal pool of electrodes.

**Table 1.** Mean correlations between connectivity fluctuation and frontal oscillatory power. The table lists the Spearman correlations between fluctuation amplitude and the frontal power, averaged across an established frequency band (e.g., Delta is the average of 1 Hz power, 1.5 Hz power, … 3.5 Hz power). The effects for Delta, Theta, Beta, and Gamma remain significant if significance testing is instead performed using permutation-testing and with family-wise error rate correction (p_corrected_ < .05). The significance stars represent uncorrected significance levels from a standard t-test. *, p < .05; **, p < .01; *** p < .0001.

| Frequency ranges | | Mean correlation | Cohen's $d$ |
| --- | --- | --- | --- |
| Delta | 1-3.5 Hz | .042 [.027, .056] | 1.24 *** |
| Theta | 4-7.5 Hz | .041 [.025, .056] | 1.13 *** |
| Alpha | 8-12.5 Hz | .029 [.012, .047] | 0.71 ** |
| Beta | 13-29.5 Hz | .029 [-.002, .059] | 0.41 * |
| Gamma | 30-50 Hz | .037 [.010, .064] | 0.59 * |

Similar albeit weaker trends emerged when correlating fluctuation amplitude with power averaged across a larger pool of electrodes, including central and parietal ones (**Figure 7B**). Confirmatory analyses demonstrated that these oscillatory links specifically reflect associations with the posterior-anterior/ventral-dorsal absolute difference and not the signed difference or the overall level of connectivity (Supplemental Materials 6). These patterns are most consistent with a characteristic timescale near 3–6 Hz for the amplitude of the putative high/low-PE fluctuations. This is notably consistent with established links between PE and Delta/Theta^30,39^ and is further consistent with an interpretation in which these fluctuations relate to PE-related processing during rest.

## 3. Discussion

The present research conducted task-fMRI, rs-fMRI, and rs-fMRI-EEG studies to clarify whether PE elicits global connectivity effects and whether the signatures of PE processing arise spontaneously during rest. This investigation carries implications for how PE minimization may characterize abstract task-general cognitive processes. *First*, the two task-fMRI studies (Studies 1A/1B) yielded converging evidence that high-PE stimuli upregulate connectivity between ventral and dorsal regions (OC-IPL and ATL-LPFC), whereas low-PE stimuli upregulate posterior-anterior connectivity (OC-ATL and IPL-LPFC). *Second*, the two rs-fMRI studies (Studies 2A/2B) indicated that the brain, even while at rest, shifts between global states of strong posterior-anterior connectivity (low-PE signature) and strong ventral-dorsal connectivity (high-PE signature). Importantly, the recruitment of ventral-dorsal and posterior-anterior networks during rest correlated with the recruitment of these networks during a PE-processing task across participants. *Finally*, the rs-fMRI-EEG study (Study 3) showed that these connectivity fluctuations correlate primarily with Delta/Theta oscillatory (EEG) power. Although there are different ways to interpret this correlation, it is consistent with high/low PE states generally fluctuating at 3-6 Hz during rest. Below, we discuss these three studies’ findings.

The connectivity patterns put forth are, for the most part, not spatially novel and instead overlap heavily with prior functional and anatomical findings. Detecting that high PE upregulates ventral-dorsal (OC-IPL & ATL-LPFC) connectivity matches the handful of prior studies on how PE influences interactions between regions. On the posterior side of the ventral-dorsal network, finding that high PE enhanced OC-IPL connectivity is consistent with PE prompting attentional shifts by the parietal cortex.^8,9^ For this interpretation and those upcoming, note that these mechanisms will likely only apply to subsets of the full ROIs; the OC and IPL are functionally heterogeneous structures, and our analyses focused more on broad-scale patterns rather than drawing neuroanatomically narrow conclusions. On the anterior side of the ventral-dorsal network, the association between high PE and ATL-LPFC connectivity aligns with findings on PE signals flowing between ATL and PFC.^42,43^ Given ATL’s role in encoding semantics and abstract information,^13,44–46^ ATL-LPFC interactions could reflect conflicts between incoming stimuli and existing mental models detected in ATL and propagated to PFC,^43^ and/or signals from the PFC to update categories in ATL.^43^ Altogether, ventral-dorsal connectivity could underlie interactions between temporal lobe representations and frontoparietal control processes, with OC-IPL connectivity reflecting more perceptual interactions and ATL-LPFC connectivity representing more semantic interactions.

In contrast with high PE, low PE was tied to greater connectivity between posterior and anterior brain regions. On the ventral side of the posterior-anterior network, low PE enhanced OC-ATL connectivity along the ventral stream. This finding is consistent with perception studies showing how the ventral stream encodes stimulus predictions (schemas) that guide processing.^47^ Strong OC-ATL connectivity under low PE could reflect information automatically flowing along the ventral pathway, in the absence of the frontoparietal control processes demanded by high PE. On the dorsal side of the posterior-anterior network, low PE boosted IPL-LPFC connectivity, which points to the relevancy of the frontoparietal network, although this network boasts many functions and its exact role for low PE processing is unclear.^48^ Overall, these posterior-anterior patterns recapitulate well-known dorsal frontoparietal and ventral stream pathways that have been central to the focus of attention and visual processing, respectively.

These functional pathways, which form the backbone of our PE topology, also recapitulate respective structural dorsal (superior longitudinal fasciculus; SLF) and ventral (inferior longitudinal fasciculus, ILF, uncinate fasciculus) pathways. Supporting these inferences, the SLF has been associated with cognitive control and attention,^49,50^ and the ILF has been associated with the semantic processing of objects,^51^ faces,^52^ and words.^53^ It is therefore unsurprising that low PE is associated with such well-worn, structurally connected pathways, and future studies may help to shed light on how major fiber pathways support a global topology of prediction error. Furthermore, that high PE is associated with a connectivity pattern that runs contralaterally to these well-worn pathways is also consistent with the idea that novel connectivity patterns are necessary to form coherent associations between schematically incongruent stimuli. Nonetheless, the consistencies between functional and structural topologies serve to build an inferential bridge linking the cognitive processes at play in PE tasks to resting state.

Our rs-fMRI investigation examined whether resting dynamics resemble the task-defined connectivity signatures of high vs. low PE, independent of the type of stimulus encountered. The resting-state analyses indeed found that, even at rest, participants’ brains fluctuated between strong ventral-dorsal connectivity and strong posterior-anterior connectivity, consistent with shifts between states of high and low PE. This conclusion is based on correlative/observational evidence and so may be controversial as it relies on reverse inference; it is impossible to establish that the task networks only reflect PE and that the rs-fMRI patterns are not due to some other variable manipulated by the task. Nonetheless, several key concerns are addressed. First, we established these signatures across two distinct PE designs (Studies 1A/1B), constraining the possible interpretations of the networks. Second, we linked the recruitment of ventral-dorsal and posterior-anterior networks during rest to the engagement of these networks during a PE task in the same participants (Study 2B), creating a sharp parallel between the task effects and resting-state functioning. Although further work is warranted to confirm the function of ventral-dorsal and posterior-anterior during rest, the present research remains, we believe, the most thorough investigation of the type of abstract PE processes that lie at the core of contemporary theories.

A key finding supporting our interpretation is the significant link between individual differences in task-evoked PE responses and resting-state fluctuations. One might initially view the effect size of this correspondence (r = .021) as modest. However, this interpretation must be contextualized by the considerable measurement noise inherent in short task-fMRI scans; the split-half reliability of the task contrast was only .08. This low reliability imposes a severe statistical ceiling on any possible task-rest correlation. Therefore, detecting a highly significant (p < .001) relationship despite this constraint provides robust evidence for a genuine link. Furthermore, our analytical approach, which iteratively examined thousands of ROI subsets rather than one aggregated network, was intentionally granular. The goal was not simply to correlate two global measures, but to test for a person-specific anatomical correspondence – that is, whether the specific parts of an individual’s network most sensitive to PE during the task are the same parts that fluctuate most strongly at rest. An aggregate analysis would obscure this critical spatial specificity. Taken together, this granular analysis provides compelling evidence for an anatomically consistent fingerprint of PE processing that bridges task-evoked activity and spontaneous resting-state dynamics, strengthening our central claim.

While our claims about resting-state cognition will often be speculative, it is worth considering what minimization of PE could look like in a resting-state scan, or more succinctly during the cognitions that we expect to be active during rest.^54,55^ One possibility is that PE could compel episodic replay or episodic simulation, consistent with the notion that perceptual decoupling and episodic memory retrieval are necessary conditions for the self-generation of a train of thought unrelated to the external environment.^56,57^ For instance, in their stream of thought, a participant’s working memory may disengage from their immediate scanner environment, and retrieve information about a recent discussion with a colleague. In turn, the participant may wonder how this discussion resolved (PE detection; high PE), prompting them to retrieve a similar experience to resolve this uncertainty (PE response; low PE). This description casts PE as a spark for episodic replay.^15^ In turn, the participant may consider that their comment was rude (PE detection) and predict how their colleague will treat them in the future (PE response). To be clear, these types of mental processes are not exclusive to resting-state scans and could likewise occur when participants experience an event live. Yet, by showing resting dynamics that resemble task-defined PE signatures, we suggest that the currently identified networks are not necessarily limited to lower-level sensory violations. In turn, the present findings support theories about how many aspects of executive function and high-level cognition are rooted in PE minimization.^4–7,58^

Finally, our third approach for investigating the global PE hypothesis was to examine if high and low PE networks fluctuate sufficiently fast for cognitive operations. As noted before, executive function EEG studies suggest that PE detection^26,27^ and PE minimization^26,28,29^ are separated by a few hundred milliseconds. This result matches evidence that PE modulates Delta (1-4 Hz) and Theta (4–8 Hz) power.^30^ Thus, we expected high and low PE networks would fluctuate multiple times per second. Indeed, the fMRI-EEG correlations in Study 3 suggest that the ventral-dorsal and posterior-anterior networks fluctuate at a speed around 3-6 Hz. We specifically interpret the fMRI-EEG correlation as reflecting fluctuation speed because we correlated EEG oscillatory power with the fluctuation amplitude computed from fMRI data. Simply correlating EEG power with the average connectivity or the signed difference between posterior-anterior and ventral-dorsal connectivity yields null results (Supplemental Materials 6), suggesting that this is a very particular association, and viewing it as capturing fluctuation amplitude provides a parsimonious explanation. Yet, this correlation may be interpreted in other ways. For example, resting-state Theta is also a signature of drowsiness,^59^ which may correlate with PE processing, but perhaps should be understood as some other mechanism. Additionally, Theta is widely seen as a sign of dorsal anterior cingulate cortex activity,^39^ and it is unclear how to reconcile this with our claims about network fluctuations. Nonetheless, as we show with simulations (Supplemental Materials 5), a correlation between slow fMRI network fluctuations and fast EEG Delta/Theta oscillations is also consistent with a common global neural process oscillating rapidly and eliciting both measures.

Some prior studies have correlated EEG spectral signatures with fMRI functional connectivity in resting-state data,^60^ including by correlating fMRI measures with power, which was likewise averaged over intervals and convolved with a hemodynamic response function.^38^ However, our work was unique because we specifically correlated power with the absolute difference between connectivity measures after establishing that those measures correspond to dynamic connectivity states. In turn, our interpretation shifts. Whereas these prior papers claimed, for instance, that an inverse correlation between functional connectivity and alpha indicates ties to alertness,^38^ we put forth that the correlations with oscillations reveal the frequency of the high/low PE-state fluctuations – an interpretation supported by our simulations.

Interpreting connectome-wide phenomena as changing multiple times per second may seem unlikely, given that much network research is based on fMRI and its slow temporal resolution. However, prior studies incorporating data from other modalities have likewise argued that, despite usually being measured slowly, network states can fluctuate at sub-second frequencies.^61,62^ Theoretically, the identified association between PE topology fluctuations and power in Delta/Theta frequencies agrees with the premise that these fluctuations reflect ongoing human thought. Such thoughts can be conceptualized as repeated manipulations of working memory, which can occur up to five times per second in humans^63^ and is indexed by Delta/Theta oscillations.^64^ Further research on other variables remains necessary to draw firm conclusions about specific cognitive mechanisms, although these timelines are consistent with the types of functions discussed by psychological PE theories – e.g., working memory states prompting memory recall or affective labeling to minimize PE could conceivably occur in ∼200 ms.^4,15^

To conclude, PE elicits brain-wide effects on connectivity, upregulating ventral-dorsal connectivity while down-regulating posterior-anterior connectivity. These patterns resemble global connectivity signatures seen in resting-state participants, and correlations between fMRI and EEG data yield associations, consistent with participants fluctuating between high-PE (ventral-dorsal) and low-PE (posterior-anterior) states at 3-6 Hz. Although definitively testing these ideas is challenging, given that rs-fMRI is defined by the absence of any causal manipulations, our results provide evidence consistent with PE minimization playing a role beyond stimulus processing. These findings hence empirical foundation for the deluge of theories on the role of PE in abstract cognition. PE minimization may be the *telos* of the brain, while attentional reorienting, episodic retrieval, working memory updating, and other psychological processes may be how PE minimization is accomplished. In this way, the present results link neuroscientific literature while also joining this work with more cognitively focused theories of the mind.

## Data availability

The datasets used for Studies 1B and 2B are publicly available from the Human Connectome Project website (https://db.humanconnectome.org). The dataset for Study 3 is also publicly available (https://fcon_1000.projects.nitrc.org/indi/retro/nat_view.html). The data collected for Studies 1A and 2A will be shared upon request via email to the corresponding author, Paul C. Bogdan, or principal investigators Roberto Cabeza and Simon W. Davis. Note that the results supporting the main conclusions of Studies 1A and 2A were each replicated using the freely available Study 1B and 2B datasets.

## Code availability

The Python and R code used for every study has been made publicly available (https://github.com/paulcbogdan/FlucPE). This repository also contains the result images directly generated via the analysis code.

## Acknowledgments

This work was supported by the National Institutes of Health, R01-AG066901, R01-AG075417.

## 4. Materials and Methods

### 4.1. Study 1A – Task-fMRI

#### 4.1.1. Participants

The Study 1A task-fMRI analyses were performed using data from a larger study on semantic processing, memory, and aging. Participants were recruited from the local community and screened for no history of brain injury or mild cognitive impairment. Sixty-six healthy participants completed the PE task, which included a younger adult subset (N = 35, M_age_ = 22.8 [SD = 3.3]; sex: 64% female, 36% male) and an older adult subset (N = 31, M_age_ = 71.5 [SD = 4.5], 64% female, 36% male). The two groups were not compared here. Of the 66 participants, two encountered scanner malfunctions that caused one run of data loss from each, but these partial datasets were still included in the analyses. There were no exclusions. All participants provided informed consent, the manuscript includes no information that could be used to identify participants, and the research was approved by the Duke University Health System Institutional Review Board.

#### 4.1.2. Task design

Participants viewed pictures of common scenes and objects. Each trial began with a picture of a scene (e.g., a farm or desert) shown for 3 seconds. Then, a blank screen appeared for an average of 3 seconds (jittered), followed by a picture of an object shown for 4 seconds. While the object was displayed, participants used a 4-point scale to indicate how likely it would be to find the object in the scene (1 = “*Very unlikely*”, 4 = “*Very likely*”). Across three runs, participants completed 114 trials: 38 high PE, 38 medium PE, and 38 low PE. For the high-PE condition, the object-scene pairs were incongruent – e.g., a *poker table* (object) likely would not be found in a *prison* (scene) because prisoners’ leisure is restricted. For the medium PE condition, the object and scene were meant to be conceptually orthogonal – e.g., a *bug* could be found in a *desert* but not necessarily. For the low PE condition, the pairs were congruent – e.g., a *tractor* operates on a *farm.* All participants saw the same 114 objects and 114 scenes but counterbalanced under different configurations. For instance, the *tractor* was low PE for one-third of participants, medium PE for one-third, and high PE for one-third. The Results response times based on the 77.2% of trials where participants gave the expected response: 1 likelihood for high PE and 4 likelihood for low PE. Of note, the medium PE condition yielded slower responses than both conditions (M = 2.33 s, SD = 0.43) (comparisons of high/low to medium show a large difference, *p*s < .0001, *d*s > 1.96). These response time patterns suggest that the medium condition was more difficult, but low vs. high differences in the brain data would not be attributable to difficulty. An analysis that treats PE as a gradient from low-medium-high would be well positioned to establish that effects genuinely reflect PE rather than task difficulty, which would instead be reflected by strong medium PE connectivity. As part of the aforementioned wider memory study, on a previous day, participants completed a task where they viewed the objects without proceeding scenes. Additionally, on the day following the PE task, participants completed a surprise memory retrieval test. However, neither of these tasks’ data were analyzed here.

#### 4.1.3. MRI acquisition

MRI data were collected using a General Electric 3T MR750 scanner and an 8-channel head coil. Anatomical images were acquired using a T1-weighted echo-planar sequence (96 slices at 0.9 × 0.9 × 1.9 mm^3^). Functional images were acquired using an echo-planar imaging sequence (repetition time = 2000 ms, echo time = 30 ms, field of view = 19.2 cm, 36 oblique slices with voxel dimensions of 3 × 3 × 3 mm). Stimuli were projected onto a mirror at the back of the scanner bore, and responses were recorded using a four-button fiber-optic response box (Current Designs, Philadelphia, PA, USA). Functional resting-state images were collected from the participants using the same parameters (210 volumes, 7 minutes).

The below descriptions of anatomical and functional preprocessing were automatically generated by fMRIPrep with the express intention that users should copy and paste this text into their manuscripts unchanged.

#### 4.1.4. Anatomical data preprocessing

A total of two T1-weighted (T1w) images were found within the input BIDS dataset. All of them were corrected for intensity non-uniformity (INU) with N4BiasFieldCorrection,^65^ distributed with ANTs 2.3.3.^66^ The T1w-reference was then skull-stripped with a Nipype implementation of the antsBrainExtraction.sh workflow (from ANTs), using OASIS30ANTs as a target template. Brain tissue segmentation of cerebrospinal fluid (CSF), white-matter (WM), and gray-matter (GM) was performed on the brain-extracted T1w using fast (FSL 6.0.5.1).^67^ An anatomical T1w-reference map was computed after registration of two T1w images (after INU-correction) using mri_robust_template (FreeSurfer 7.3.2).^68^

Brain surfaces were reconstructed using recon-all (FreeSurfer 7.3.2),^69^ and the brain mask estimated previously was refined with a custom variation of the method to reconcile ANTs-derived and FreeSurfer-derived segmentations of the cortical gray-matter of Mindboggle.^70^ Volume-based spatial normalization to one standard space (MNI152NLin2009cAsym) was performed through nonlinear registration with antsRegistration (ANTs 2.3.3), using brain-extracted versions of both T1w reference and the T1w template. The following template was selected for spatial normalization and accessed with TemplateFlow (23.0.0):^71^ ICBM 152 Nonlinear Asymmetrical template version 2009c.^72^

#### 4.1.5. Functional data preprocessing

For each of the seven BOLD runs found per subject (across all tasks and sessions), the following preprocessing was performed. [The mention of “seven” BOLD runs here reflects other tasks collected as part of the aforementioned memory study.] First, a reference volume and its skull-stripped version were generated using a custom methodology of fMRIPrep. Head-motion parameters with respect to the BOLD reference (transformation matrices, and six corresponding rotation and translation parameters) are estimated before any spatiotemporal filtering using mcflirt (FSL 6.0.5.1, Jenkinson et al. 2002). BOLD runs were slice-time corrected to 0.972s (0.5 of slice acquisition range 0s-1.94s) using 3dTshift from AFNI.^73^

The BOLD time series (including slice-timing correction when applied) were resampled onto their original, native space by applying the transforms to correct for head motion. These resampled BOLD time-series will be referred to as preprocessed BOLD in original space, or just preprocessed BOLD. The BOLD reference was then co-registered to the T1w reference using bbregister (FreeSurfer) which implements boundary-based registration.^74^ Co-registration was configured with six degrees of freedom.

Several confounding time series were calculated based on the preprocessed BOLD: framewise displacement (FD), DVARS, and three region-wise global signals. FD was computed using two formulations following Power et al. (absolute sum of relative motions)^75^ and Jenkinson (relative root mean square displacement between affines).^76^ FD and DVARS are calculated for each functional run, both using their implementations in Nipype (following the definitions by Power et al.^75^). The three global signals are extracted within the CSF, the WM, and the whole-brain masks. Principal components are estimated after high-pass filtering the preprocessed BOLD time series (using a discrete cosine filter with 128s cut-off) for the two CompCor variants: temporal (tCompCor) and anatomical (aCompCor). tCompCor components are then calculated from the top 2% variable voxels within the brain mask. For aCompCor, three probabilistic masks (CSF, WM, and combined CSF+WM) are generated in anatomical space. The implementation differs from that of Behzadi et al. in that instead of eroding the masks by two pixels on BOLD space, a mask of pixels that likely contain a volume fraction of GM is subtracted from the aCompCor masks. This mask is obtained by dilating a GM mask extracted from the FreeSurfer’s aseg segmentation, and it ensures components are not extracted from voxels containing a minimal fraction of GM. Finally, these masks are resampled into BOLD space and binarized by thresholding at 0.99 (as in the original implementation). Components are also calculated separately within the WM and CSF masks. For each CompCor decomposition, the k components with the largest singular values are retained, such that the retained components’ time series are sufficient to explain 50 percent of variance across the nuisance mask (CSF, WM, combined, or temporal). The remaining components are dropped from consideration.

The head-motion estimates calculated in the correction step were also placed within the corresponding confounds file. The confound time series derived from head motion estimates and global signals were expanded with the inclusion of temporal derivatives and quadratic terms for each.^77^ Additional nuisance time series are calculated by means of principal components analysis of the signal found within a thin band (crown) of voxels around the edge of the brain, as proposed by Patriat, Reynolds, and Birn.^78^ The BOLD time series were resampled into standard space, generating a preprocessed BOLD run in MNI152NLin2009cAsym space. First, a reference volume and its skull-stripped version were generated using a custom methodology of fMRIPrep. All resamplings can be performed with a single interpolation step by composing all the pertinent transformations (i.e. head-motion transform matrices, susceptibility distortion correction when available, and co-registrations to anatomical and output spaces). Gridded (volumetric) resamplings were performed using antsApplyTransforms (ANTs), configured with Lanczos interpolation to minimize the smoothing effects of other kernels.^79^ Non-gridded (surface) resamplings were performed using mri_vol2surf (FreeSurfer). Many internal operations of fMRIPrep use Nilearn 0.9.1,^80^ mostly within the functional processing workflow. For more details of the pipeline, see the section corresponding to workflows in fMRIPrep’s documentation.

#### 4.1.6. Functional connectivity

Calculating ROI-ROI functional connectivity required first measuring each voxel’s BOLD response in each trial. This was done using first-level general linear models with the Least Squares Separate approach by Mumford et al.,^81^ which involves fitting a separate regression for each trial. The regression included a boxcar signal spanning the trial’s object presentation period, and this regressor’s coefficient represents the trial’s BOLD response. The regression also included a boxcar signal covering the presentation time of all other stimuli (i.e., every other object and every scene). Additionally, the linear models included six translation/rotation regressors, three other covariates for head motion (FD, DVARS, and RMSD), and covariates for mean global, white-matter, and cerebrospinal signals.

The voxelwise measures of activity (beta coefficients) were organized and averaged in terms of the 210 neocortical ROIs of the Brainnetome Atlas (i.e., the cortex sans the hippocampus); the exact ROIs employed for the main analyses involving four anatomical quadrants are shown in Supplemental Figure S1. ROI-ROI functional connectivity was calculated independently for the high/medium/low PE conditions as the Pearson correlation between two vectors describing trial-wise activation estimates for each pair of ROIs (beta-series correlations).^82^ Notably, the Brainnetome Atlas provides anatomical labels for each ROI. However, it does not provide an “ATL” label, which was defined here as the eight most anterior temporal gyral ROIs, as our group has previously done in semantic processing research.^45^ The data on ROI-ROI connectivity was used for the modularity analysis (described next) and for the classification-based analyses, which evaluated connectivity effects while assessing statistical significance with multiple-comparison correction (methods and results provided in Supplemental Materials 2). The analyses were also conducted and reproduced using the Schaefer atlas (Supplemental Materials 7).

#### 4.1.7. Modularity

Our goal in testing modularity and our approach contrasts typical modularity studies, which identify task-independent patterns by examining the connectivity averaged across conditions and participants. Here, modularity was tested using a beta matrix representing the effect of PE on each connection. For each edge, a regression was fit predicting connectivity strength, while treating PE as a continuous predictor (low = -1, medium = 0, high = +1). The regression was analogous to a paired t-test but with three points rather than two. That is, each participant contributed one sample per PE condition, and the regression included a dummy variable for each participant (66 participants = 65 subject variables). This was separately done for each edge in the connectome to construct the aforementioned beta matrix. Evaluating modularity required preparing binary matrices. Specifically, the beta matrix was filtered to only the 5% of edges with the most positive coefficients (high > low PE) or the 5% of edges with the most negative coefficients (low > high PE). The high-filtered and low-filtered binary matrices were then submitted to the Leiden algorithm – an optimized form of Louvain’s algorithm, which addresses the risk of disconnected communities and has become increasingly recommended.^83^

### 4.2. Study 1B – Task-fMRI

#### 4.2.1. Participants and task design

Analyses were conducted on 1,000 participant’s gambling task data from the S1200 HCP dataset (M_age_ = 28.7, 57% female, 43% male); this dataset is described by Barch et al.^34^ The task is based on the design by Delgado et al.^84^ with some changes to the timing procedures.^34,85^ In each trial, participants were shown a question mark and asked to press a button to make a choice (“2” or “3”) within 1.5 s. After making a choice, participants immediately saw a fixation cross for the rest of the 1.5 s period. Then, the fixation continued for another 1 s. Afterward, participants saw an upward green arrow in 43.75% of trials (signifying a $1.00 reward) or a downward red arrow in 43.75% of trials (signifying a $0.50 loss). In 12.5% of trials, participants encountered a screen indicating a neutral outcome (no gain or loss). Participants were told that their choice determined the outcome, but the choice actually had no effect, and each trial’s outcome was predetermined. The monetary difference between rewards/losses was meant to account for loss aversion causing losses to be perceived as worse than gains. Participants completed two runs of the task, each with 32 trials separated into four blocks.

#### 4.2.2. Task design and modeling PE

To model PE, analyses defined a participant’s prediction in each trial as an exponential moving average, *EMA[t] = α × outcome + (1 – α) × EMA[t – 1]*. Outcomes were defined as +1, losses as -1, and neutral outcomes as 0. The learning rate was defined at 0.3, although testing 0.2 or 0.5 showed that this does not change whether the Connectivity Direction × PE interaction effect is significant (still *p* < .0001). The prediction at the start of each run was defined as zero. PE was computed as the absolute difference between a trial’s outcome and the prediction based on previous trials.

This continuous measure of PE was converted into binary high-PE and low-PE conditions to support the functional connectivity analyses. Binarization involved dividing trials based on the median PE. This median split was done separately for gain and loss trials to ensure they were in equal quantities across the high/low PE conditions. The neutral trials were omitted from the connectivity analysis as they were few in number. The first trial of the run was also omitted as there were no prior trials on which to base a meaningful prediction. Although a median split will lead to roughly equal numbers of trials in the high/low PE conditions, to ensure that these two had exactly the same number of trials (rather than possibly one condition having one or two more), trials at the end of the task were omitted for balancing as needed. To validate these conditions and the binary split, participants’ reaction times were compared post-high-PE-loss and post-low-PE-loss trials using a paired t-test, which was expected to show post-error slowing.^31^

#### 4.2.3. Imaging protocol and preprocessing

Details on the imaging protocols are provided by Van Essen et al.^33^ Of note, the functional voxels were small (2 mm isotropic) and the temporal resolution was very fast (0.72 s TR time). The downloaded dataset had already been “minimally preprocessed” by the HCP, which included the removal of spatial artifacts/distortions, segmentation, within-subject registration, and normalization to a common brain template.^86^ Although the imaging and preprocessing protocols differ in Study 1B from Study 1A, this is acceptable given that the goal for Study 1B was not to be a precise replication. Additionally, adhering to the already-performed preprocessing for a public dataset benefits reproducibility.

#### 4.2.4. Functional connectivity

Analysis of functional connectivity paralleled that of Study 1A but with two changes. First, single-trial beta estimates were computed for each trial using a Least Squares All rather than a Least Squares Separate approach.^81^ The Study 1B dataset is much larger than the Study 1A dataset, and the Least Squares All approach is much less computationally intensive; preliminary analyses suggested that the produced beta values are largely similar. Second, each single-trial-beta regression model included the motion parameters provided by the HCP (six translation/rotation regressors and their derivatives) along with a covariate for mean global signal; note that the Study 1A preprocessing did not include motion derivatives, but this was done here because the HCP provided them, and we sought consistency with their standard procedures. Functional connectivity estimates were calculated separately for high-PE and low-PE conditions, for each pair among the 210 ROIs by using beta-series connectivity, as above.^82^ The contrast matrix was correlated with that of Study 1A (**Figure 2A**). The Results section reports the correlation taken between the bottom triangle of a 46x46 matrix, wherein ROIs were averaged based on their Brainnetome label (e.g., averaging all L MFG ROIs to a single region); this is the matrix shown in **Figures 2A & 3A**. This averaging yields a more stable measure, given that individual signal within ROIs can vary wildly due to noise. As reported, the correlation was fairly strong (*r* = .27, *p* < .0001). If the correlation is instead taken with respect to the 210x210 matrix, there remains a clear correlation albeit a weaker one (*r* = .14, *p* < .0001).

### 4.3. Study 2A – Resting-state-fMRI

#### 4.3.1. Data and preprocessing

Rs-fMRI data were collected from the 66 participants from Study 1A with the same acquisition protocols, and also as above, with the six motion covariates regressed out. However, whereas the single-trial models for the task-fMRI analysis included a global-signal covariate, this was not done for the rs-fMRI analysis because global signal regression would strongly bias ROI-ROI and edge-edge correlations toward negative values.^87,88^ This would be an issue for the rs-fMRI analyses, as they focused on pure correlations rather than contrasts between conditions. Yet, without global signal regression, connectivity is strongly biased toward positive correlations.^89^ Hence, our resting-state analyses analyzed only a subset of the brain (OC, ATL, IPL, LPFC) and removed effects linked to connectivity with regions outside these subsets. This was done by data cleaning with temporal CompCor (see Section 4.1.5) and with other procedures detailed in the next subsection.

#### 4.3.2. Edge-edge correlations

Our analysis of the rs-fMRI data focused on how connectivity strength fluctuates over time and how groups of edges fluctuate in sync or out of sync. Of the emerging techniques for addressing dynamic connectivity,^90^ we chose to employ *time-varying connectivity* and *edge-edge correlations*.^35,91,92^ Whereas traditional connectivity studies have focused on the correlation between activation time series extracted from pairs of ROIs, this direction models connectivity itself as a time series and quantifies the relations between edges’ time series. Time-varying connectivity is modeled by decomposing the traditional Pearson correlation.^35,91,92^ That is, a Pearson correlation entails z-standardizing two vectors and taking the average of their element-wise products. For time-varying connectivity, the element-wise products are calculated but not averaged, which means that the two input time series have created a new time series of the same length. For example, in the first volume, if OC activity is high and IPL connectivity is high, the OC-IPL connectivity in the first volume is said to be strong. After time-varying connectivity has been calculated for each edge in the connectome, the values were averaged in terms of the major anatomical quadrants shown in **Figure 3A**. For example, the OC-IPL entry represents the average across every possible pair of OC and IPL ROIs.

In turn, Pearson correlations can be computed between pairs of the edge sets (e.g., correlating OC-IPL × OC-ATL). This edge-edge correlation represents how the two edges shift over time in sync (positive correlation) or out of sync (negative correlation). As mentioned in the prior subsection, these analyses required accounting for global signal. Preliminary tests showed that if global signal effects are unaccounted, this would promote a very positive correlation between edges, particularly for pairs of edges linked to the same ROI (i.e., global signal particularly enhances correlations between ROI_1_-ROI_2_ × ROI_1_-ROI_3_). Yet, if traditional global signal regression was done during preprocessing, this would induce a very negative correlation. Hence, to achieve an unbiased estimate, rather than measuring edge-edge connectivity as plain Pearson correlations, they were measured as partial correlations. These partial correlations essentially measured edge-edge fluctuations relative to each edge’s baseline correlation with the rest of the connectome. For this, analyses gathered the 144 neocortical ROIs outside the four quadrants (non-OC, -IPL, -ATL, or -LPFC) and calculated the mean time-varying connectivity between each octant with these 144 ROIs (e.g., the mean time-varying connectivity between the L OC and the rest of the brain) (a octant is the unilateral version of a quadrant; see **Figure 4**). These eight measures of time-varying connectivity, one with each octant, were then partialled out of each edge-edge correlation. This partial correlation strategy is analogous to existing methods for unbiased global signal regression designed for classic ROI-ROI connectivity, which regress out signal from regions not analyzed. This approach appeared to be successful, given that in the correlation matrix shown in **Figure 4**, the mean correlation is *r* = .02, suggesting the absence of pressure toward positive or negative correlations.

Edge-edge correlations were measured between each of the twelve connections shown in the **Figure 4** cuboid to produce an edge-edge correlation matrix, on which modularity was tested. In this context, modules constitute sets of edges that are all strong in the same volumes (positive edge-edge correlations within modules). Modularity also is based on fluctuation between modules (negative correlations between modules). For the modularity analysis, the variant of Louvain’s algorithm was used as above but now, rather than just filtering the matrix to the 5% of strongest edges, the matrix was filtered to the 50% of strongest edges. This much more liberal approach was necessary as the edge-edge correlation matrix was much smaller (i.e., for a 12×12 matrix, using just 5% of edges would mean that most rows do not have a single preserved entry).

### 4.4. Study 2B – Resting-state-fMRI

#### 4.4.1. Data and preprocessing

Rs-fMRI data were used from the 1,000 HCP participants of Study 1B, which were collected using the same acquisition protocols and had already been submitted to the HCP minimal preprocessing pipeline. Preprocessing was meant to be similar to that of Study 2A, including regressing out the six motion parameters and using temporal CompCor but not global signal regression. Afterward, the rs-fMRI data were submitted to the edge-edge modularity analysis described above for Study 2A, attempting to reproduce the original modules

#### 4.4.2. Analysis

Study 2B also incorporated a new analysis, considering subsets of the four anatomical quadrants (visualized in **Figure 5**). Specifically, the analysis was predicated on repeatedly selecting a pair of bilateral ROIs (one left and its symmetric right) at random from the four quadrants. This selection yields eight ROIs, and the strength of the Study 1B task-fMRI effect and the strength of the rs-fMRI fluctuation were computed for each participant. The task-fMRI effect constituted the Connectivity Direction × PE interaction, (PA_High PE_ – PA_Low PE_) – (VD_High PE_ – VD_Low PE_); these values refer to a single participant’s posterior-anterior (PA) and ventral-dorsal (VD) effects for a given condition. The rs-fMRI effect was computed using time time-varying (see above, Section 4.3.2), and the effect was defined as the mean of the TR-by-TR absolute difference between posterior-anterior and ventral-dorsal connectivity, E[|z(PA) – z(VD)|]; posterior-anterior and ventral-dorsal connectivity were each z-scored to ensure that the measure does not merely reflect overall variability in participants’ BOLD data. This absolute difference measure would later be used and discussed more prominently in Study 3, and the simulations (detailed in full in Supplemental Materials 5) demonstrate and provide intuition for why this quantity captures the degree of fluctuation between two states in resting-state fMRI data.

The task-fMRI and rs-fMRI measures were computed for each participant and for each selection of eight ROIs (3,432 in total). They were then submitted to correlations and group-level analyses. Specifically, within-ROI-set (subject-by-subject) correlation was computed for each set of ROIs, and the correlation coefficients were likewise submitted to a one-sample t-test assessing whether the mean correlation significantly surpassed zero. In addition, a multilevel regression was performed which accounted for subjects’ overall task-resting association with respect to the full quadrants – i.e., to show that the results indeed reflect within-subject heterogeneity in the anatomy of their PE networks rather than just their intensity. All 3.432 million data points (3,432 sets × 1,000 participants) were submitted to the following multilevel regression: *rest ∼ 1 + task + (1 + task | ROI set) + (1 | subject)*. The use of random slopes for the ROI-set-level grouping makes this analogous to the t-test done on the correlations for each ROI set. The results report the significance of the *task* fixed effect, computed using the *lmerTest* R package.^93^

#### 4.4.3. Links between network fluctuations and behavior

We considered whether the extent of PE-related network expression states during resting-state is behaviorally relevant. We specifically investigated whether individual differences in the overall magnitude of resting-state fluctuations could predict individual difference measures, provided with the HCP dataset. This yielded a significant association with age, whereby older participants tended to display weaker fluctuations. However, associations with cognitive measures were limited. A full description of these analyses is provided in Supplemental Materials 8.

### 4.5. Study 3 – Rs-fMRI-EEG

#### 4.5.1. Participants and preprocessing

The fMRI-EEG analyses used the public dataset provided by Telesford and colleagues^37^, which consists of fMRI and EEG data simultaneously recorded from 22 participants (M_age_ = 36.8 [22-51], 50% male, although one participant’s data was unusable, see below). Participants generally completed two 10-minute resting-state scans. Participants also completed two 10-minute scans where they viewed the *Inscapes* video – an animation featuring shapes that slowly move and transform, which is meant to mirror resting-state conditions while discouraging movement and drowsiness.^94^ Data for one resting-state scan and one Inscapes scan were missing from five participants. Also, in our analyses, we encountered difficulties temporally aligning ten scans’ fMRI data and EEG data. Hence, the analyses were done using 68 scans, spread across 21 participants. The dataset also includes scans where participants viewed movies with higher cognitive demands (e.g., Hollywood films), but these were not used. For ease of reproducibility, the analyses used the preprocessed version of the dataset provided by Telesford et al.^37^; see their report for a description of the acquisition and preprocessing procedures. Of note, the voxel size was 3.469 x 3.469 x 3.330 mm and the TR length was 2100 ms. The preprocessing by Telesford et al. notably included gradient artifact removal, which introduced discontinuities to the EEG data. For the fMRI data, we further preprocessed the data using temporal CompCor, as otherwise, as was done for the resting-state Studies 2A and 2B and as without CompCor, the edges all strongly correlated.

#### 4.5.2. Time-frequency analysis

The time-frequency analyses were done using electrode data recorded at 5000 Hz, which we downsampled to 250 Hz. The analyses first targeted five midline frontal electrodes (F3, F1, Fz, F2, F4; BioSemi64 layout), given that this frontal row is typically the focus of executive-function PE research on oscillations.^30,40,41^ Second, given the large size of the posterior-anterior/ventral-dorsal fMRI pattern, analyses were also done with the frontal and the frontocentral (FC3, FC1, FCz, FC2, FC4), central (C3, C1, Cz, C2, C4), centroparietal (CP3, CP1, CPz, CP2, CP4), and parietal (P3, P1, Pz, P2, P4) electrodes. As part of their preprocessing, Telesford et al.^37^ omitted some electrodes’ data (different across participants, M = 3.2 electrodes omitted). For the time-frequency analysis, complex Morlet seven-cycle wavelets were defined for each frequency band 0.5-50 Hz in 100 linearly spaced 0.5 Hz steps. Each electrode’s time series was convolved using the wavelets, and the resulting spectrograms were averaged across electrodes for each participant. The discontinuities in the data generated during preprocessing, as mentioned above, create border artifacts in the time-frequency data, and the next section describes how these were dealt with.

#### 4.5.3. Fluctuation-power correlation

Using the fMRI data, time-varying connectivity was calculated for each edge among the quadrant pairs illustrated earlier (e.g., edges between each of the OC ROIs with each of the ATL ROIs). Connectivity was averaged by quadrant pair as above, yielding overall time-varying measurements for OC-ATL, IPL-LPFC, OC-IPL, and ATL-LPFC connectivity. Partialling out edge-edge correlations with the rest of the brain, as done for the rs-fMRI studies, was not necessary here, as our focus was not on comparing positive vs. negative correlations while computing modularity. At each TR, the absolute difference (|PA – VD|) was calculated between posterior-anterior connectivity (OC-ATL + IPL-LPFC) and ventral-dorsal connectivity (OC-IPL + ATL-LPFC). This time series of absolute differences represents how the intensity of the posterior-anterior vs. ventral-dorsal fluctuations shifts over time. That is, fluctuation amplitude was considered equivalently high in TRs when (i) posterior-anterior connectivity was strong while ventral-dorsal was weak or (ii) ventral-dorsal connectivity was strong while posterior-anterior was weak. This measure is discussed in Supplemental Materials 5, and simulations are provided showing how this measure captures the amplitude of an underlying neural oscillator, even if the oscillator’s frequency outpaces fMRI’s temporal resolution. This absolute difference was correlated with oscillatory power at different frequency bands. The simulations in Supplemental Materials 5 also show why measuring such correlations shed light on the frequency underlying the fMRI connectivity fluctuations.

In the EEG data, the measures of oscillatory power were organized by TR and averaged. For instance, at the fourth volume, the oscillatory power of a given frequency was averaged across 6300-8399 ms. That is, the spectrograms’ temporal resolutions were reduced from 250 Hz to 1/2.1 Hz. To account for EEG discontinuities introduced during preprocessing, and the potential for discontinuities to create border artifacts, TR intervals overlapping with discontinuous segments were identified along with their neighbors (e.g., if TR 4 overlaps with a discontinuous 500 ms segment, then TRs 3-5 are marked). Oscillatory power in these TRs was interpolated linearly (e.g., for TRs 3-5, interpolated using TRs 2 and 6). Finally, before oscillatory power was correlated with the connectivity-fluctuation differences, the interpolated spectrograms were convolved with respect to time using a hemodynamic response function. This accounts for the lag in the fMRI data between neural activation and the later BOLD response. After organization, the analysis simply consisted of measuring the Spearman correlation between the connectivity-fluctuation difference and oscillatory power at each frequency bin (e.g., correlate 1 Hz, 1.5 Hz, etc.). For each participant, this correlation was measured for each of their four scans, then Fisher z-transformed, and averaged. Each participant’s mean correlation was submitted to a one-sample t-test assessing whether the correlation significantly surpassed zero.

To confirm the significance of our fMRI-EEG correlations with a non-parametric approach, we performed a group-level permutation-test. For each of 1,000 permutations, we phase-randomized the fMRI fluctuation amplitude time series. Specifically, we randomized the Fourier phases of the |PA–VD| series (within run), while retaining the original amplitude spectrum; inverse transforms yielded real surrogates with the same power spectrum. This procedure breaks the true temporal relationship between the fMRI and EEG data while preserving its structure. We then re-computed the mean Spearman correlation for each frequency band using this phase-randomized data. We evaluated significance using a family-wise error correction approach that accounts for us analyzing five oscillatory power bands. We thus create a null distribution composed of the *maximum* correlation value observed across all frequency bands from each permutation. Our observed correlations were then tested for significance against this distribution of maximums.

