## Supplemental Materials for "Spontaneous fluctuations in global connectivity reflect transitions between states of high and low prediction error"

### 1. Exact ROIs used

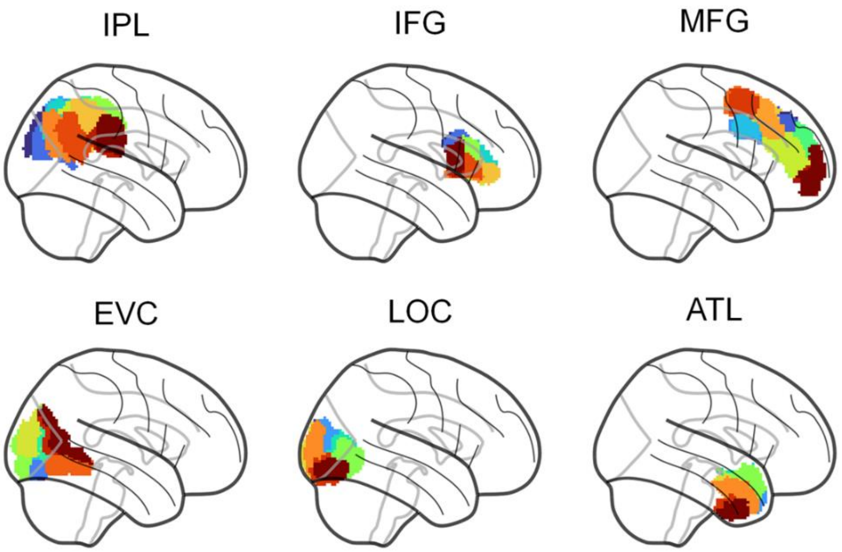


***Figure S1. Illustration of the ROIs used***. *The areas used for the four quadrants are shown here. For each area (IPL, IFG, etc.), the different colors distinguish among the different Brainnetome ROIs under this label assigned by the atlas.*

### 2. Evaluating the statistical significance of the Study 1A effects

#### 2.1. Network-Based Statistic

Here, we evaluate whether PE significantly impacts connectivity at the network level using the Network-Based Statistic (NBS) approach.[7] NBS relied on the same regression data generated for the main-text analysis, whereby a regression is performed examining the effect of PE (Low = –1, Medium = 0, High = +1) on connectivity for each edge. This was done across the connectome, and for each edge, a z-score was computed. For NBS, we thresholded edges to $|Z| > 3.0$, which yielded one large network cluster, shown in Figure S3. The size of the cluster – i.e., number of edges – was significant ($p < .05$) per a permutation-test using 1,000 random shuffles of the condition data for each participant, as is standard.[7] These results demonstrate that the network-level effects of PE on connectivity are significant. The main-text modularity analysis converts this large cluster into four modules, which are more interpretable and open the door to further analyses.

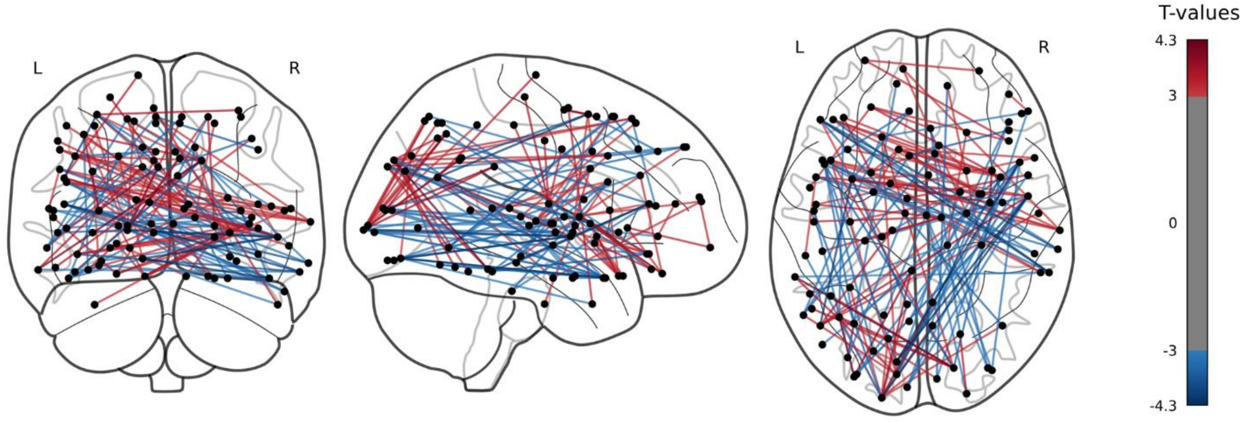


**Figure S3. Network-Based Statistic results for Study 1A.** Visual examination of the cluster roughly points to the same four posterior-anterior and ventral-dorsal modules identified formally in main-text Figure 2C.

#### 2.2. ROI-based classification analysis

To identify the effects of prediction error (PE) on functional connectivity, for Study 1A, we began with a classifier-based approach, given the high statistical power that this affords.[1–3] The analysis targeted the 210 neocortical ROIs from the Brainnetome Atlas. For each ROI, a support vector machine (SVM) was fit, making binary predictions and attempting to distinguish the high- and low-PE conditions. For a given ROI, its classifier used all of the connections to said ROI as features (**Figure S2**). The input dataset was organized such that each participant contributed two examples/observations to each ROI's analysis: one high-PE example and one low-PE example. The examples' edges were mean-centered within-subject by subtracting the edge's average activity across the subject's two examples; this makes the analysis analogous to a paired t-test. Altogether, the 66 participants' data produced 132 examples. Each ROI's SVM accuracy was assessed via group-2-fold cross-validation, averaged across 100 repetitions; see work on why this cross-validation configuration maximizes sensitivity.[4]

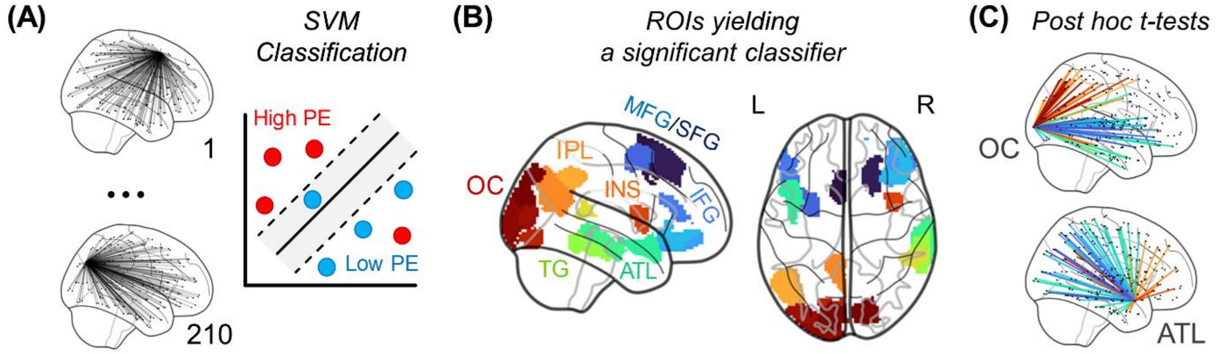


***Figure S2. Classifier-based analysis on the connectivity effects of prediction error. (A)*** *210 linear support vector machines (SVMs) were trained and tested, one for each ROI. The SVMs attempted to label examples as high or low prediction error.* ***(B)*** *Every ROI that yielded significant classifier accuracy is shown here, colored arbitrarily.* ***(C)*** *Post hoc-tests are shown based on the connections to occipital cortex (OC) and*

*anterior temporal lobe (ATL) ROIs that yielded a significant classifier. Hot colors indicate stronger high-PE connectivity and cool colors indicate stronger low-PE connectivity. SFG, superior frontal gyrus; MFG, middle frontal gyrus; IFG, inferior frontal gyrus; INS, insula; TG, temporal gyri (inferior/middle/superior).*

The statistical significance of each classifier's testing accuracy was evaluated via permutation testing. For the permutations, the examples were shuffled within-subject to preserve the dataset's intrinsic structure.[5] Because each participant contributed two examples, shuffling amounted to randomly selecting participants to have their low-PE and high-PE labels flipped. For the permutation tests, the examples were submitted to the exact analyses described above, which included iteration across every ROI. This was done 100 times, and given that there were 210 ROIs, permutation testing. yielded 21,000 measures of testing accuracy. The resulting distribution was used to assign each ROI's classifier an uncorrected p-value (e.g., if the ROI achieved 99.5th percentile accuracy relative to the 21,000 estimates, it was assigned $p_{uncorrected} = .005$). The p-values were then corrected in terms of a false-discovery rate.[6] Significance ($p_{FDR} < .05$) required at least 60% accuracy.

Functional connectivity analysis followed the procedures described above. Several ROIs' classifiers yielded above-chance accuracy, and these ROIs are shown in **Figure S1B**. In other words, these regions' functional connections with the rest of the brain were modulated by PE. This classifier-based analysis establishes the statistical significance of PE-related connectivity changes linked to these brain areas. Yet, the nature of machine learning approaches requires post hoc analyses to draw clear neuroscientific interpretations. **Figure S1C** shows the beginning of this, presenting the results of paired t-tests associated with connections to occipital cortex (OC) and anterior temporal lobe (ATL) ROIs that each yielded a significant classifier However, the primary interpretation tests consist of the modularity analysis in the main text portion of Study 1A, and this analysis also serves to establish the PE signatures used for the rs-fMRI studies.

### 3. BOLD activation contrasts

#### *3.1. Study 1A*

The effect of PE on single-region BOLD signal was examined to establish a more complete description of the brain data. This was conducted using an ROI-based approach, analogous to that used for the main text connectivity analysis. For Study 1A (semantic PE), each participant's BOLD signal in each ROI was averaged for each of the three conditions (low/medium/high PE). Each ROI was then analyzed independently using a linear regression that predicted BOLD based on a PE predictor (low = -1, medium = 0, high = +1). The regression included a dummy variable representing each subject's identity, meaning that the analysis is akin to a paired t-test. This strategy produced the contrast shown in **Figure S3A**. Prominently, low PE upregulates lateral parietal and lateral temporal activity. High PE upregulates medial occipital, precuneus, and ATL activity. Several of these emerging areas overlap with the regions linked to functional connectivity changes in the main text, which is consistent with these regions being indeed involved in PE processing. Note that this data does not see any obvious signs that the main text conclusions on connectivity derive from effects that can be explained in terms of activity confined within individual regions (i.e., coactivation rather than connectivity).[1] Rather, the main text PE connectivity results seem to indeed be best understood as PE influencing the relationship between regions. Indeed, when the main text regression of connectivity strength on Connectivity-Direction x PE interaction is performed while including each quadrant's mean activity as a covariate (four covariates), the original interaction effect remains significant ($\beta = .14$, $p < .001$).

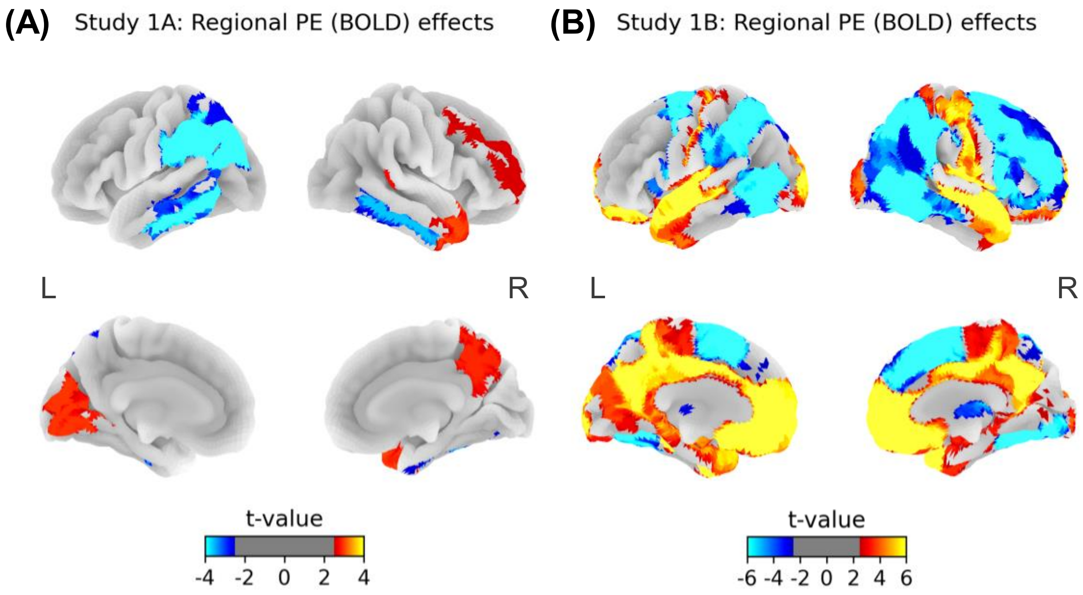


***Figure S3. Effects of prediction error on regional activation.*** ***(A)*** *These t-maps correspond to the Study 1A analysis of PE's effect on BOLD. Note that a liberal threshold is used (p < .05 [two-tailed]), as the focus is on being descriptive rather than drawing specific conclusions. The ROIs employed here use the Brainnetome atlas, as for the main text analysis, but to increase power at the expense of anatomical precision, ROIs were grouped based on their area label and averaged (e.g., the L IPL ROI's signal here is based on the average of the six L IPL ROIs).* ***(B)*** *These t-maps correspond to the Study 1B analysis of PE's effect on BOLD. Unlike for the Study 1A analysis, given that Study 1B had more participants, all 210 neocortical Brainnetome ROIs were analyzed independently without area averaging.*

#### *3.2. Study 1B*

BOLD analyses were also performed for Study 1B (gambling PE task), again based on ROIs but now simply submitting the low/high PE conditions' data to a paired t-test as there are just two conditions (**Figure S3B**). Each of the patterns associated with Study 1A emerged again. However, there are also some new effects. In particular, the ventromedial prefrontal cortex stands out showing a strong link to high PE. This is presumably due to the region's central role in processing outcomes and win/loss values.[8] Additionally, although these regions are not clearly

[1] It is also not clear whether connectivity "artifacts" induced by BOLD coactivation would even be mathematically possible. For papers focusing on just pairs of regions (e.g., the ATL and OC), then both regions' BOLD rising in one condition can sometimes cause the condition to also be seen as increasing connectivity. However, we do not see any way that isolated BOLD effects could produce the four-quadrant pattern producing diverging posterior-anterior or ventral-dorsal effects.

visible in **Figure S3B**, the effects of high > low PE are large in the ventral caudate ($t$[999] = 10.92, $p < .0001$), dorsal caudate ($t$[999] = 9.74, $p < .0001$), and ventral striatum ($t$[999] = 8.92, $p < .0001$); analyses combining the left and right ROIs. Overall, the Study 1A results being largely consistent with Study 1B but also partially divergent speaks to these two tasks engaging common PE mechanisms while also being complementary.

**4. Within-subject consistency of PE networks and fluctuations**

The main text poses that, even at rest, participants' brains are characterized by fluctuations between low PE and high PE network states. If this indeed reflects meaningful resting-state phenomena, then these network states and the fluctuations should be stable within individuals such that their neural processing is consistent across scans. This is particularly relevant to the Study 2B analyses, given that they are predicated on modeling heterogeneity in these networks across individuals. For Study 2B, participants completed each resting-state scan twice, and the present analyses focus on linking the pairs of scans from each participant. We evaluated stability in two ways: (i) whether within-subject heterogeneity in static functional connectivity is consistent across separate resting-state scans and (ii) whether within-subject heterogeneity in dynamic connectivity fluctuation is also consistent across separate resting-state scans.

For the first analysis, we focused on stability in terms of the static properties of the presented networks – by "static," we simply refer to a classic measure of functional connectivity across a full scan as the Pearson correlation between two ROI's BOLD time series. We computed each participant's functional connectome for each of their two resting-state scans. Then, we isolated the edges among one pair of adjacent quadrants (e.g., just IPL-LPFC edges). The consistency of the networks' patterns across scans was first measured via within-subject edge-by-edge correlations between the edge strengths across the two scans.[2] This was done for each participant, then averaged across participants. For all four pairs of quadrants, consistency was substantial (OC-IPL $r = .72$; ATL-LPFC $r = .62$; IPL-LPFC $r = .83$; OC-ATL $r = .43$). Similar patterns emerge if we perform the analogous across-subject consistency analysis – i.e., for each edge, examining whether participants with high connectivity strength in one scan tend to also display high strength in another scan (mean correlations: OC-IPL $r = .51$; ATL-LPFC $r = .52$; IPL-LPFC $r = .62$; OC-ATL $r = .35$). To be clear, these results do not speak directly to the question of whether these resting-state patterns reflect PE, which is covered elsewhere. However, the results demonstrate the stability of functional connectivity measurements generally.

For the second analysis, we focused on whether the dynamic properties of participants' networks (the fluctuation patterns) are also stable across scans. Our analysis paralleled the conceptualization used in the main text where posterior-anterior and ventral-dorsal network states fluctuate between one another. These analyses are also similar to those of earlier work on edge-edge correlations.[9] We computed time-varying connectivity for each participant's connectome. Then, we focused on pairs of edges from adjacent quadrants associated with one overlapping ROI

[2] This is equivalent to measuring the (within-subject, ROI-wise) statistical reliability of a measurement. The second set of values reported lower in the paragraph represent the across-subject statistical reliability.

(e.g., one OC-ATL edge and one ATL-LPFC edge); analogous to how an edge was the minimum "unit" for analysis of the above stability analysis, here this minimum unit is a pair of edges with one common ROI.[9] For a given participant and edge pair, the magnitude of fluctuation between the two edges was computed as in the main text: namely, as the mean absolute difference of two edges' time-varying connectivity time series. For each participant, fluctuation magnitudes are computed for all possible sets of three ROIs among an anatomical quadrant triplet. Then, the within-subject triplet-by-triplet correlation is computed between a participant's two scans, which is averaged across participants. As in the analysis of static connectivity, we find that fluctuation magnitude is a stable quantity across resting state scans (OC-*IPL*-LPFC $r = .53$; IPL-*LPFC*-ATL $r = .65$; LPFC-*ATL*-OC $r = .53$; ATL-*OC*-IPL $r = .65$). Likewise, there is high consistency in the analogous across-subject analysis like above (OC-*IPL*-LPFC $r = .44$; IPL-*LPFC*-ATL $r = .42$; LPFC-*ATL*-OC $r = .46$; ATL-*OC*-IPL $r = .39$). Thus, the network fluctuations targeted are stable and reflect reliable individual differences.

**5. Simulating the effects of oscillations on fMRI and EEG data**

The third study's fMRI-EEG analyses are based on two premises: (1) Fluctuations between posterior-anterior (PA) and ventral-dorsal (VD) global states could produce the identified fMRI patterns, even if the oscillations outpace fMRI's temporal resolution. (2) By correlating the PA vs. VD absolute differences with EEG oscillatory power at multiple frequencies, this can shed light on the frequency of the PA/VD fluctuations. To validate these premises and provide intuition for them, a simulation study was performed.

This simulation conceptualizes the constant transitions between high-PE (ventral-dorsal) and low-PE (posterior-anterior) states as deriving from an oscillating mechanism (the sine wave in **Figure S4A**). The two premises above can be derived from the idea that this oscillator's amplitude varies over time, and this variation changes more slowly than the oscillator's frequency. For instance, suppose that a participant shifts between high-PE and low-PE states at 3 Hz. If the participant feels anxious or thinks about something important for 4 seconds, this may cause high-amplitude 3 Hz oscillations for 4 seconds. To simulate these types of relatively slow changes, amplitude can be defined in terms of an autoregressive process (shown in **Figure S4B**):

$$A(t+1) = |r * A(t) + \sqrt{1 - r^2} * \sigma|$$

$\sigma$ represents noise sampled from a normal distribution, drawn independently at each time point. The equation takes the absolute value of the autoregressive process because amplitude cannot be negative. As shown in **Figure S4B**, this process shifts over time, and the high amplitude period from 5-9 seconds can be interpreted as representing the acute 4-second period of anxiety mentioned above. Together, the product of the sine wave and amplitude produces an oscillator with varying intensity over time (**Figure S4C**):

$$x(t) = A(t) * \sin(ft)$$

$f$ is the oscillator's frequency ($f$ = 3 Hz for **Figure S4**). As this oscillator is meant to represent shifts between primarily PA or VD connectivity, high values of $x(t)$ correspond to strong PA connectivity (low-PE state) and low values of $x(t)$ correspond to strong VD connectivity (high-PE state) (or vice versa). Based on this signal, measured fMRI data (**Figures S4D-F**) and EEG time-frequency data can be simulated (**Figures S4G-I**), as will be described.

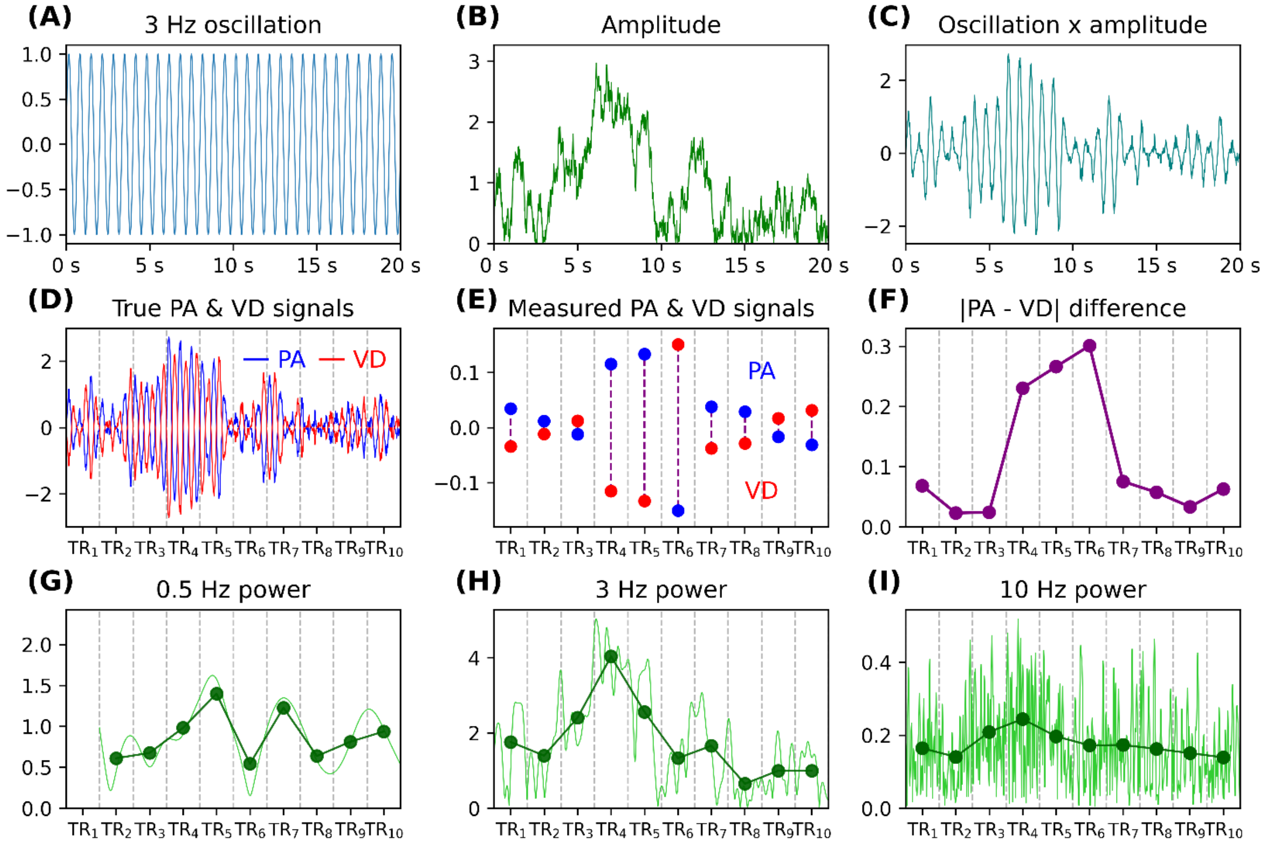


***Figure S4. Quantities used by the simulation. (A)** One sine wave is used as an input for the simulation. **(B)** An autoregressive time series is also used as an input. **(C)** The product of the sine wave and autoregressive time series defined the simulated oscillator, which represents how fluctuations between states may vary in intensity over time. **(D)** Ventral-dorsal (VD) connectivity is defined as the sine-amplitude product, and posterior-anterior (PA) connectivity is defined as the inverse of this product. **(E)** Representing brain data recorded using fMRI, the VD and PA dots here are the average over the TR windows. No hemodynamic response function convolution was applied when making this figure for the sake of clarity but convolution was performed for the simulation results reported in the text. **(F)** Using the VD and PA dots, the |VD – PA| differences were computed. **(G-I)** The green lines represent the output of time-frequency decomposition done using the sine-amplitude product. The dots represent the power averaged for each TR window. The first TR window for 0.5 Hz power was removed as it was susceptible to edge artifacts.*

PA connectivity at each moment in time can be represented as simply the oscillator itself ($x$), and VD connectivity can be represented as the inverse of the oscillator ($-x$) (**Figure S4D**). In turn, the PA and VD measurements recorded using fMRI can be computed by averaging the data over 2 s windows and convolving with the hemodynamic response function (HRF) (the dots in **Figure S4E** represent the fMRI measurements at each 2 s TR):

$$\mathrm{PA}(t') = h(1 - x'(t') + \sigma)$$

$$\mathrm{VD}(t') = h(x'(t') + \sigma)$$

The apostrophe ($'$) is used to represent averaging/down-sampling to 2 s resolution. HRF convolution is represented using $h$. Using these quantities, the PA and VD absolute difference can be computed as in the main text (**Figure S4F**):

$$d(t') = |PA(t') - VD(t')|$$

For the EEG analysis, the oscillator is decomposed into time-frequency data, representing oscillatory power at different frequencies using Morlet wavelets as described in the main text.

$$\mathrm{W}_f(t) = M(x(t) + \sigma)$$

$M$ represents transforming the oscillator signal into frequencies using Morlet wavelets. $\mathrm{W}_f$ represents the power time series recorded at a given frequency ($f$). **Figures S4G-I** show the power time series computed for 0.5 Hz, 3 Hz, and 10 Hz power. Notice how the green line for 3 Hz power best mimics the initial amplitude series because the oscillator's true frequency is 3 Hz. The subsequent steps describe how this can be derived from the data with fMRI-EEG correlations.

To link the EEG data with the fMRI measurements, the two must be first aligned in terms of temporal resolution and hemodynamic effects must be accounted for. Hence, the time-frequency data was averaged in terms of 2 s windows and convolved with the HRF:

$$\mathrm{W}''_f(t') = h(\mathrm{W}'_f(t'))$$

**Figures S4G-H** represent this 2-second window averaging using the dark green dots (representing the average) above the light green lines (representing the moment-by-moment power time series).

Then, the correlation between the PA-VD difference ($d$) and oscillatory power ($\mathrm{W}''$) can be computed for different frequencies. One would expect that the correlation between the PA and VD absolute difference ($d$) with oscillatory power ($\mathrm{W}''$) is strongest for the frequency that matches the one used to generate the oscillation, which is 3 Hz for the present example. Accordingly, in the visualization, notice how the PA and VD difference (**Figure S4F**) bears similarity to the 3 Hz power data (**Figure S4H**), evidenced by the $TR_4$ and $TR_5$ dots being elevated for both.

To show this quantitatively, data was simulated using the above equations with the following parameters: 2 s TR for the fMRI data, sampling resolution of 100 Hz for the EEG data, amplitude autocorrelation of $r = .95$, and a true oscillation frequency of 3 Hz. The degree of noise was set to be fairly large ($\sigma$ was drawn from a normal distribution with a standard deviation of 0.5). The length of the simulated time series was very long (100,000 seconds) to ensure that estimates are reliable, and to account for this simulation effectively just being of a single person whereas an actual study would examine multiple, as was indeed done for Study 3.

Correlating neural data, fMRI measurements, and EEG measurements yields patterns supporting the initial two premises: (1) The PA and VD absolute difference, $d(t')$, was correlated with the true oscillator amplitude, $A(t')$, which yielded a positive association ($r = .028$, $p < .0001$).

Hence, a sub-second oscillator will influence fMRI data even though fMRI's temporal resolution is slower than the oscillation frequency. (2) The PA and VD absolute difference was positively correlated with 3 Hz power ($r$ = .052), and this correlation was stronger than with power at any other frequency, 1 such as 1 Hz ($r$ = .005), 5 Hz ($r$ = .01 Hz), or 10 Hz ($r$ = .003). Note that this correlation peaking primarily with the power of the true frequency occurs regardless of the frequency level. For instance, when the true oscillation is set to 1, 10, or 30 Hz, the strongest correlations emerge between the PA and VD absolute difference and 1, 10, or 30 Hz power, respectively.

Finally, note that for the sake of visualization, Figure 7A shows fMRI measures (PA and VD) sampled from the peaks and troughs of the oscillations. However, in reality, the fMRI data would sample from any point in the oscillation's phase (e.g., sometimes near the midpoint, where there are virtually no PA and VD differences), and for frequencies above 0.5 Hz, fMRI would induce averaging. However, as the present simulations demonstrate, this does not invalidate the analyses but rather would just introduce noise. This may explain why the correlations are numerically small (e.g., $r$ = .052 above and the correlations reported in Table 1 of the main text).

**6. Confirmatory analyses on the link between fluctuations and oscillations**

To ensure that the Study 3 correlations between oscillatory power and |PA - VD| indeed reflect connectivity fluctuations rather than overall strong or weak connectivity, we reperformed the Study 3 analysis while adding three alternative connectivity measures as covariates: (i) the signed difference between posterior-anterior and ventral-dorsal connectivity, (ii) the sum of posterior-anterior and ventral-dorsal connectivity, and (iii) the absolute value of the sum of posterior-anterior and ventral-dorsal connectivity. The within-subject Spearman correlations were replaced with linear regression of oscillatory power on fluctuation amplitude alongside the three covariates. The links between |PA - VD| and Delta/Theta remained significant (Delta: $t[20] = 3.45$, $p = .003$; Theta: $t[20] = 5.12$, $p < .0001$). Hence, these patterns confirm that the EEG oscillations were specifically tied to connectivity fluctuation amplitude.

We additionally tested the relationship between each of those three measures and the five EEG oscillation bands. Across the 15 tests, there were no associations ($ps_{\text{uncorrected}} \geq .04$); one uncorrected p-value was at $p = .044$, although this was expected given that there were 15 tests. Thus, the association between EEG oscillations and the fMRI measure is specific to the absolute difference (i.e., amplitude) measure.

**7. Schaefer atlas analysis**

The main-text analyses all use the Brainnetome atlas, but for the sake of robustness and generalizability, the analyses were also performed using the Schaefer atlas (400-ROI default variant).[10] Because the Schaefer atlas defines its own area labels that differ from those of the Brainnetome atlas, we first needed to establish which Schaefer areas best correspond to those of the four quadrants based on the Brainnetome atlas. This was done by identifying which Schaefer areas had a majority of voxels overlapping with one of the four quadrants. For instance, in the Schaefer atlas, there exists a "L PFCl" label, and 88% of L PFCl voxels overlap with the

Brainnetome L MFG or L IFG voxels. Thus, for the Schaefer analysis, L PFCl ROIs were used for the dorsal anterior quadrant. This strategy was used to develop a four-quadrant layout using the Schaefer ROIs, which was in turn used to reproduce the findings from the different main-text analyses.

#### *7.1. Study 1*

Analyses were first done on the semantic PE data (Study 1A). Computing connectome-wide PE regressions and modularity analysis again yields the two high PE and two low PE modules as before (**Figure S5A**; note that this is the one analysis that does not rely on the anatomically defined quadrants, here all 400 ROIs were used to define the connectome matrix). In addition, the Connectivity Direction x PE interaction remains significant (*β [standardized] = .14, p* = .009). Next, analyses were performed on the gambling PE data (Study 1B), and again, the Connectivity Direction x PE interaction remains significant ($F[1, 999] = 8.47$, $p = .003$).

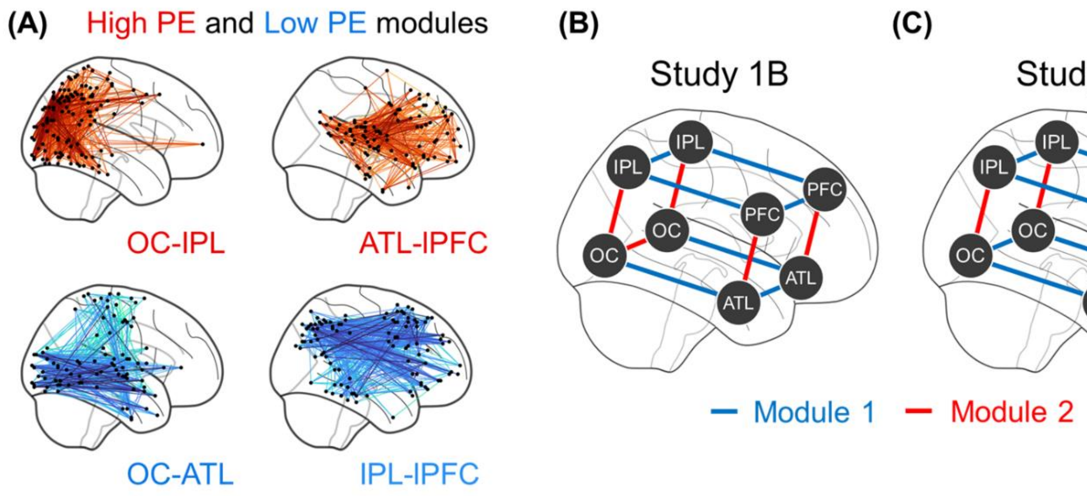


***Figure S4. Replication using the Schaefer atlas.*** *These figures correspond to main text* ***(A)*** *Figure 2C,* ***(B)*** *Figure 4C, and* ***(C)*** *Figure 5.*

#### *7.2. Study 2*

Analyses next reproduced the key elements of the dynamic connectivity modules shown in Studies 2A and 2B (**Figures S5B & S5C**). There are notably some differences in which modules the interhemispheric connections are assigned to, but the overall split across posterior-anterior and ventral-dorsal dynamic connectivity states remains the same. The original task-fMRI × rs-fMRI analysis for Study 2B was not possible for the Schaefer atlas given as the atlas is not bilaterally symmetric, precluding the selection of eight ROI sets.

#### *7.3. Study 3*

Finally, fMRI-EEG analyses reproduced the patterns initially reported in main-text Table 1, where the strongest association between (fMRI) connectivity fluctuation magnitude and EEG oscillatory power is seen with Delta and Theta (Table S1)

***Table S1. Mean correlations between connectivity fluctuation and frontal oscillatory power.*** *The table lists the Spearman correlations between fluctuation amplitude and the frontal power, averaged across an established frequency band (e.g., Delta is the average of 1 Hz power, 1.5 Hz power, ... 3.5 Hz power). This is a reproduction of main-text Table 1 but based on using the Schaefer atlas to define the four quadrants. *, p < .05; **, p < .01; *** p < .0001.*

| Frequency ranges | | Mean correlation | Cohen's *d* |
|---|---|---|---|
| Delta | 1-3.5 Hz | .051 [.033, .070] | 1.20 *** |
| Theta | 4-7.5 Hz | .043 [.026, .059] | 1.12 *** |
| Alpha | 8-12.5 Hz | .020 [.001, .039] | 0.46 * |
| Beta | 13-29.5 Hz | .029 [.010, .048] | 0.65 * |
| Gamma | 30-50 Hz | .037 [.016, .057] | 0.77 ** |

### 8. Behavioral differences related to fluctuation amplitude

To investigate whether individual differences in the magnitude of resting-state PE-state fluctuations predict general cognitive abilities, we correlated our resting-state fluctuation measure with the cognitive and demographic variables provided in the HCP dataset.

#### *8.1. Methods*

For each of the 1,000 participants, we calculated a single fluctuation amplitude score. This score was defined as the average absolute difference between the time-varying posterior-anterior (PA) and ventral-dorsal (VD) connectivity during the resting-state fMRI scan (the average of the TR-by-TR measure used for Study 3). We then computed the Spearman correlation between this score and each of the approximately 200 individual difference measures provided in the HCP dataset. We corrected for multiple comparisons using the False Discovery Rate (FDR) approach.

#### *8.2. Results*

The correlations revealed a robust negative association between fluctuation amplitude and age, indicating that older participants tended to display weaker fluctuations ($r$ = -.16, $p_{corrected}$ < .001). After correction, two significant correlations with cognitive performance emerged: (i) a positive association with the age-adjusted score on the Picture Sequence Memory Test ($r$ = .12, $p_{corrected}$ = .03), (ii) a negative association with performance on the Card Sort Task ($r$ = -.12, $p_{corrected}$ = .046). As greater fluctuation amplitude is linked to better performance on one task but worse performance on another, it is unclear how to interpret these findings.
